# Ionizing radiation responses appear incidental to desiccation responses in the bdelloid rotifer *Adineta vaga*

**DOI:** 10.1101/2023.06.16.545282

**Authors:** Victoria C. Moris, Lucie Bruneau, Jérémy Berthe, Anne-Catherine Heuskin, Sébastien Penninckx, Sylvia Ritter, Uli Weber, Marco Durante, Etienne G. J. Danchin, Boris Hespeels, Karine Van Doninck

## Abstract

**Background:** The remarkable resistance to ionizing radiation found in anhydrobiotic organisms, such as some bacteria, tardigrades, and bdelloid rotifers has been hypothesized to be incidental to the desiccation resistance. Both stresses produce reactive oxygen species and cause damage to DNA and other macromolecules. However, this hypothesis has only been investigated in a few species.

**Results:** In this study, we analyzed the transcriptomic response of the bdelloid rotifer *Adineta vaga* to desiccation and to low- (X-rays) and high- (Fe) LET radiation to highlight the molecular and genetic mechanisms triggered by both stresses. We identified numerous genes encoding antioxidants, but also chaperones, that are constitutively highly expressed, which may contribute to the protection of proteins against oxidative stress during desiccation and ionizing radiation. We also detected a transcriptomic response common to desiccation and ionizing radiation with the over-expression of genes mainly involved in DNA repair and protein modifications but also genes with unknown functions being bdelloid-specific. A distinct transcriptomic response specific to rehydration was also found, with the over-expression of genes mainly encoding Late Embryogenesis Abundant proteins, specific Heat Shock Proteins, and glucose repressive proteins.

**Conclusions:** These results suggest that the extreme resistance of bdelloid rotifers to radiation might indeed be a consequence of their capacity to resist complete desiccation. This study paves the way to functional genetic experiments on *A. vaga* targeting promising candidate proteins playing central roles in radiation and desiccation resistance.

## Background

Living in extreme environments requires effective mechanisms of cellular and molecular protection and of repair. One of the most severe abiotic stresses for cells is desiccation. Yet, some organisms, within distinct clades such as bacteria [1], plants [2] but also metazoans, including the tardigrades [3], nematodes [4], insect larvae [5, 6], and bdelloid rotifers [7, 8] are capable of anhydrobiosis and survive extreme water loss entering a dormant or ametabolic state. Anhydrobiotic organisms were also shown to be able to survive high doses of ionizing radiation (IR). According to ***Mattimore and Battista*** [9], resistance to IR is incidental to desiccation resistance since both stresses are known to cause similar damages, including DNA double strand breaks (DSBs) [9–12]. However, this hypothesis has been supported by only limited data from a few species where they also showed that similar genes or proteins were expressed after both stresses, such as in the bacterium *Deinococcus radiodurans* [13] and in the larva of the anhydrobiotic insect *Polypedilum vanderplanki* [14]. In this study, we used the bdelloid rotifer *Adineta vaga* to shed light on the genes and biological processes activated by both stresses, desiccation and IR.

Since their discovery (***van Leeuwenhoek 1702***), bdelloid rotifers have intrigued scientists because of their capacity to survive complete desiccation at any stage of their life-cycle and their resistance to a variety of other stresses including freezing, exposure to deep vacuum or high doses of IR (i.e., >500 Gy)[10, 15–19]. Moreover, these metazoans are among the smallest animals on Earth, eutelic (i.e., a fixed number of somatic cells at maturity) and being notorious for their parthenogenetic mode of reproduction in the absence of males while having diversified into more than 460 morphospecies [20–23]. Their ability to re-start a new population from one individual and to survive complete desiccation or freezing, enables these organisms to inhabit environments where few animals can thrive, including semi-terrestrial habitats subjected to frequent episodes of drought [24] such as mosses, lichens, rain gutters, but also soil, deserts and the arctic [24–29]. Among the highly radiation and desiccation-resistant bdelloid rotifer species, *Adineta vaga* has been the most extensively studied [12, 19, 23, 30–33] and its genome has been sequenced and assembled at a chromosome-scale resolution level [34].

Both desiccation and IR are known to be associated with cellular membrane damage and the generation of Reactive Oxygen Species (ROS), inducing lipid peroxidation and DNA and protein damage [1, 35, 36]. Survival to desiccation and IR therefore requires adjustments that maintain the function of macromolecules (i.e., proteins and DNA) despite the production of ROS [37]. This can be achieved through two nonexclusive ways: either by preserving the integrity of these molecules or by repairing the damage accumulated. The loss of DNA integrity through DNA double strand breaks (DSBs) was shown to be repaired in *A. vaga* within 24-48 hours after desiccation or IR [12, 19, 23] while the study by [30] highlighted an efficient antioxidant system in *A. vaga* protecting proteins from oxidative damage. Moreover, genome analyses showed a wide expansion of gene families involved in oxidative stress resistance and DNA repair [12, 30, 31, 33, 34]. Genes coding for proteins involved in the six main DNA repair pathways conserved for the most part from prokaryotes to eukaryotes [38–41] have indeed been found in the *A. vaga* genome and transcriptome: Homologous Recombination (HR) and Non-Homologous End-Joining (NHEJ) repairing DSBs, Base Excision Repair (BER), Nucleotide Excision Repair (NER), Mismatch Repair (MR), and Alternative Excision Repair (AER) known to correct damaged or mismatched bases [31, 33]. While the expression of DNA repair genes has been studied in *A. vaga* during and post desiccation [33], their expression pattern following high doses of radiation remains to be characterized. Moreover, some proteins highly expressed in *A. vaga* following exposure to IR seem to have been acquired horizontally (***Nicolas et al., in press***). The genome of bdelloid rotifers indeed contains the highest proportion of genes of non-metazoan origin within the studied animal genomes [31, 34, 42], many being expressed following desiccation contributing to the metabolism of bdelloid rotifers [43].

Even though some damages and responses, including oxidative stress resistance and DNA repair, are hypothesized to be shared between desiccation and radiation, others might be specific to each stress factor. Indeed, anhydrobiosis, is a complex phenomenon that requires adaptations at the molecular and cellular level to face severe constraints associated with the loss of water (*i.e.*, <10% of dried mass) and causing the complete cessation of any biological activity [12, 44, 45]. Desiccation in bdelloid rotifers also induces a change of their whole body which adopts a “tun” shape, via muscle contraction, allowing an optimal resistance to stress [46]. Desiccation resistance has been associated, in plants, bacteria, fungi, extremophilic bacteria and some tardigrades, with the accumulation of bioprotectants like non-reducing disaccharides (e.g., trehalose) that vitrify the cellular matrix [3, 47]. Trehalose has not been found in bdelloid rotifers, only the trehalase gene [32, 48, 49]. Many carbohydrate enzymes were however detected in the bdelloid genomes, some being acquired horizontally [31]. It was suggested that bdelloid rotifers might protect themselves against desiccation or osmotic stress by promoting the synthesis of Late Embryogenesis Abundant (LEA) proteins or Heat Shock Proteins (HSP), also found in plants and tardigrades, believed to act as molecular shields [49–53].

IR resistance might also involve distinct, additional responses which can differ according to the type of radiation being distinguished by their Linear Energy Transfer (LET). LET is the average energy (in keV) delivered locally to the traversed matter by a charged particle over a distance of 1 μm [54–56]. Different types of IR are known to induce distinct ionization patterns. Sparse ionization (low-LET such as X-rays) produces a uniform pattern of damages throughout its target while dense ionization (high-LET, e.g., protons, Fe-ions) generates complex DNA damages including clustered DSBs [57–60]. Because these complex lesions are difficult to repair, cell killing is higher after exposure to the same dose of high-compared to low-LET radiation. The maximum effectiveness for cell killing is between 100 and 200 keV/micron in water [61]. Iron (Fe) ions at 1 GeV/n have a LET in water of 147 Kev/micron and therefore represent a highly effective radiation quality [62]. Accordingly, the survival and fertility rates were observed to be significantly reduced in *A. vaga* individuals exposed to a given dose of protons and Fe-ions in comparison with X-rays [19]. Only a few recent studies started to analyze biological processes and genes involved in responses to X-rays and charged particles [63]. However, the role of these processes and genes in bdelloid rotifer resistance and how they differ between different types of radiation has never been investigated.

Identifying the genes and processes involved in the desiccation and IR resistance of the bdelloid rotifer *A. vaga* allows us to determine the main actors of their extreme resistance and whether they are common to both stresses. We therefore analyzed here the transcriptomic response of *A. vaga* individuals exposed to desiccation and low- and high-LET radiation (respectively X-ray and Fe-ions). Our analysis allowed identifying the desiccation, post-rehydration and post-radiation core responses and highlight numerous candidate genes. Some of these genes are likely to be responsible for resistance to both stresses and could be targeted in functional studies.

## Results

### 1. Fertility of *A. vaga* decreases after exposure to high-LET radiation

We observed that longer desiccation periods (*i.e.,* 14 days) in absence of radiation were not associated with a decrease in survival and fertility in comparison to 1-day desiccated samples (***Fig S1***). Radiation also did not impact the survival rates of desiccated and irradiated individuals compared to non-irradiated controls (500 Gy X-rays: 96±3%, *vs.* 96±3% 1-day desiccated control; 500 Gy Fe: 92±2% *vs.* 91±3% GSI transport control; ***Fig S1***). However, the fertility rate after 500 Gy Fe-ions (9±5%) was lower than after 500 Gy X-ray radiation (92±7%). Indeed, 72±17% of 500 Gy Fe-ions irradiated samples were characterized by the production of sterile eggs (***Fig S1*** in blue), unable to complete full embryological development.

### 2. Transcriptomic response in *A. vaga* are highest 2.5 hours post-radiation and 1.5 hours post rehydration

With the Differential Gene Expression (DGE) analysis of *A. vaga* individuals under different treatments, including “entering desiccation” (in grey), 14 days desiccation (yellow), 500 Gy low-LET (orange or blue) and high-LET radiation (purple) (***Fig 1A***), we observed that the highest variance in gene expression (PC1=56%) correlates with the radiation treatment (irradiated vs. non-irradiated) and the time post radiation (***Fig 1B***). The maximal distance observed on PCA, is between control samples and the timepoint 2.5h after low- and high-LET radiation (***Fig 1B***). At this timepoint, we also measured a high percentage of genes under-expressed (**UE**) (in red, 6.6-13.4%) and over-expressed (**OE**) (in light green, 6.1-6.6%) with a percentage of OE genes even slightly higher at 8h following Fe-ions irradiation (8.3%) (***Fig 1C***). PC2, encompassing 16% of the variance, correlates with the desiccation treatment (desiccated vs. hydrated) (***Fig 1B***; ***Supplementary results***). Samples rehydrated for 1.5 hours (h) after 14 days of desiccation show the highest number of UE (20.4%) and OE (16.1%) genes (***Fig 1C, S2A***). We also observed that rotifers entering desiccation or post-desiccation (1.5h or 2.5h post desiccation/radiation) had more UE genes than OE genes, while we found the same percentage (6.6%) of OE and UE genes 2.5h post radiation without previous desiccation (***Fig 1C, S2B***). At 24h post-radiation, the difference between control and irradiated samples is the lowest (***Fig 1B***, pentagons) with a lower percentage of genes being differentially expressed (**DE**) (***Fig 1C***). We noticed that three hydrated controls (***Fig 1B***, yellow-green) had a positive PC2 and were more distant to the other controls. These rotifers were sampled under different food conditions (see Methods section). The replicates from the experiments with hydrated rotifers irradiated with X-rays were grouped two by two (***Fig 1B***, blue triangles, circles and squares) which is likely due to the sampling of this treatment being done on two different days.

**Fig 1.**
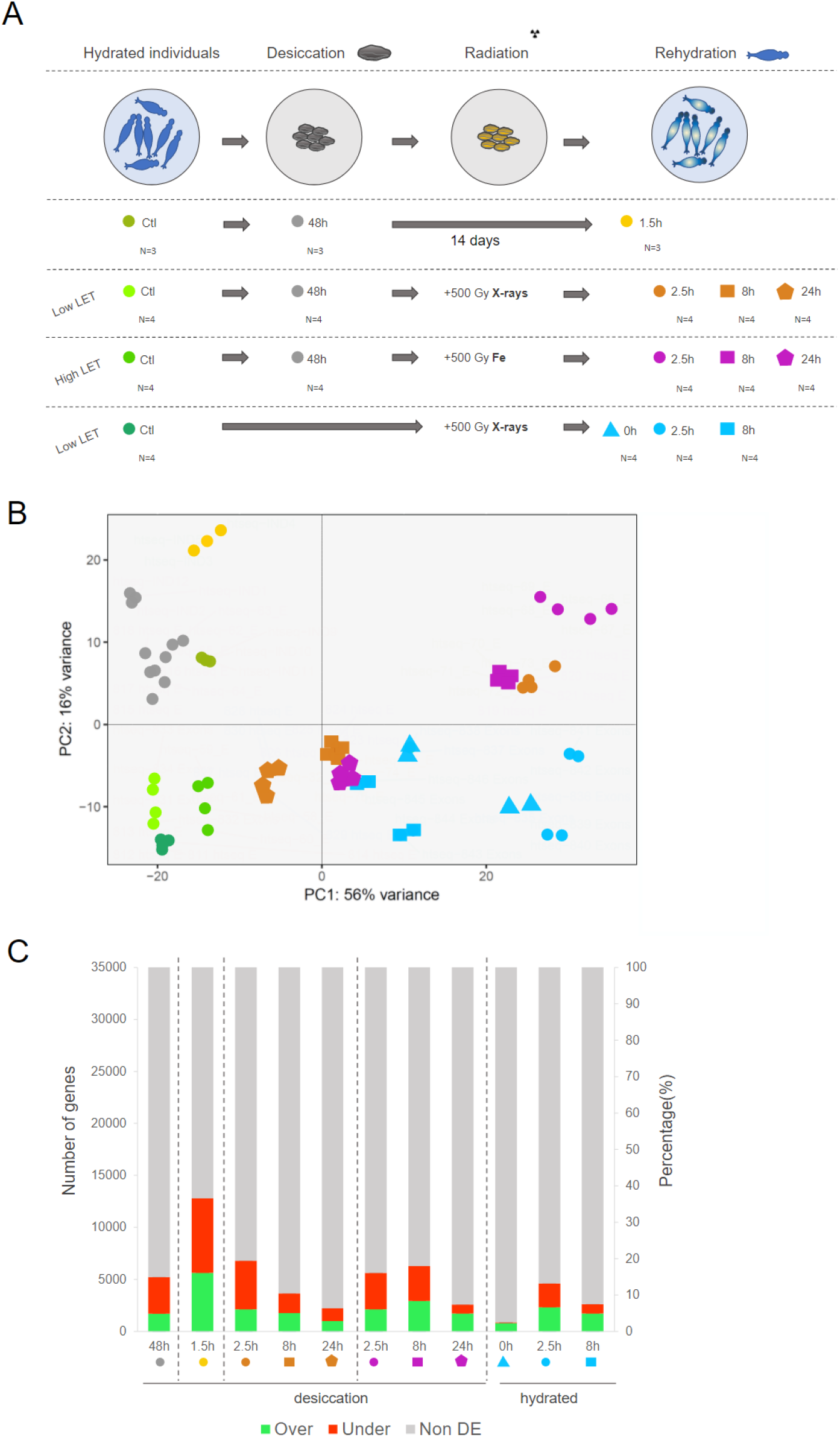
Design of the comparative transcriptomic approach and responses. A) Experimental design of the comparative transcriptomic analyses of *Adineta vaga* individuals exposed to desiccation and/or radiation: hydrated controls (in various greens), individuals entering desiccation (48 hours desiccated: in gray), rehydrated after 14 days of desiccation (in yellow), irradiated with 500 Gy X-rays (low-LET) after desiccation (in orange), irradiated with 500 Gy Fe-ions (high-LET) after desiccation (in purple), hydrated individuals irradiated with 500 Gy X-rays (in blue). The different investigated time points post-rehydration or radiation are represented as follows: 0 hours (triangles), 1.5 and 2.5 hours (circles), 8 hours (squares), 24 hours (pentagons). N = the number of replicates analyzed per condition. B) Principal component analysis of the transcriptomes analyzed (with 3 to 4 replicates per condition) with PC1 representing 56% of the variance in gene expression between samples and PC2 16% of the variance. C) Histogram of the number of genes (first y axis) and the percentage of genes within the genome (second y axis) being significantly (FDR<0.01 with both DESeq2 and EdgeR) over- (log_2_foldchange>0.5 in green) or under-expressed (log_2_foldchange<-0.5 in red) in the different conditions post-desiccation and/or radiation.

### 3. Biological processes associated to DE genes following desiccation and radiation

To identify the biological processes triggered by desiccation and by IR, GO enrichment analyses of DE genes were performed for both desiccation and radiation separately (***Supplementary Table S1***). Only half of the DE genes have GO ids attributed and could be considered for the analysis *(**Figs S3A, S3B, S3C, S3D**)*.

Some enriched biological processes (***Figs 2A***, ***2B***) and genes (***Fig 2C***) were found to be shared between *A. vaga* individuals entering desiccation (in grey) and individuals rehydrated after 14 days of desiccation (in yellow): 915 OE genes and 1885 UE genes in both conditions (***Fig 2C***, ***Supplementary Table S1***). Amongst under-expressed biological processes found significantly enriched in both conditions were processes mainly involved in signaling pathways and transport while amongst OE genes they were mostly involved in oxidation-reduction processes, ADP-ribosylation and microtubule-based processes (***Figs 2A, 2B***). Under-expressed biological processed being significantly enriched specifically to the entry in desiccation included mainly cell communication and transmembrane transport (***Fig 2A)*** while carbohydrate (e.g., glycosyl hydrolase gene families) and trehalose metabolic processes (i.e., 3 genes coding for trehalase) were enriched within OE genes (***Fig 2B***). Finally, we found significant enrichment specific to the rehydration timepoint following 14 days desiccation in UE genes involved in signaling, DNA replication and transcription, while protein modifications and signal transduction processes were enriched in OE genes (***Figs 2A, 2B***). Signaling was enriched within UE genes at the entry of desiccation and during rehydration, as well as within OE genes during rehydration suggesting some reactivation (***Figs 2A, 2B***).

**Fig 2.**
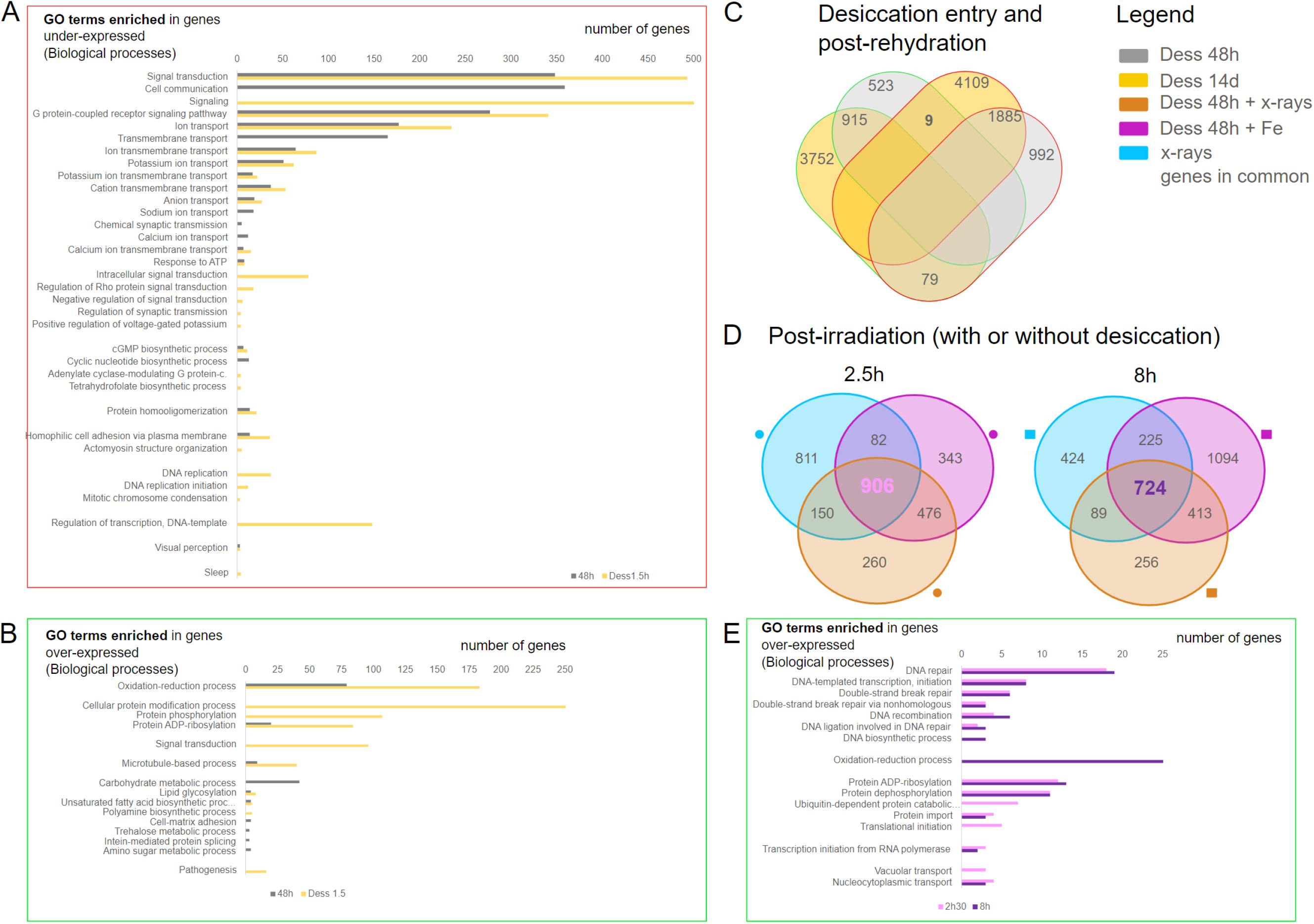
Gene Ontology enrichment analyses. A) GO biological processes significantly enriched (chi-square test p-value<0.05, min. 3 OE genes with the GO id) with genes being significantly under-expressed (framed in red, FDR<0.01 and log_2_foldchange < -0.5 in DESeq2 and EdgeR) in *A. vaga* individuals entering desiccation (in gray) and 1.5 hours post-rehydration after 14 days of desiccation (in yellow). B) GO biological processes significantly enriched (chi-square test p-value<0.05, min. 3 OE genes with the GO id) with genes being significantly over-expressed (framed in green, FDR<0.01 and log_2_foldchange > 0.5 in DESeq2 and EdgeR) in *A. vaga* individuals entering desiccation (in gray) and 1.5 hours post-rehydration after 14 days of desiccation (in yellow). C) Venn diagrams with number of genes significantly over-(green outlines) and under-expressed (red outlines) in *A. vaga* individuals entering desiccation (gray) or after rehydration and 14 days of desiccation (yellow), showing 915 genes commonly over-expressed and 1885 genes commonly under-expressed. D) Venn diagrams of genes significantly over-expressed in *A. vaga* individuals under three different conditions of radiation at 2.5h and 8h post-irradiation and/or rehydration: hydrated *A. vaga* individuals irradiated with X-rays (blue), desiccated *A. vaga* individuals irradiated with X-rays post rehydration (orange), desiccated *A. vaga* individuals irradiated with Fe post rehydration (purple). At 2.5h and 8h post-irradiation and/or rehydration, respectively 906 and 724 genes are over-expressed in the three conditions. E) GO biological processes significantly enriched (chi-square test p-value<0.05, min. 3 OE genes with the GO id) with genes found to be over-expressed in the core response to radiation (906 genes 2.5 hours post-radiation and 724 genes 8 hours post-radiation).

The same approach was applied to define the biological processes enriched in response to radiation. Similar biological processes were found to be enriched in UE genes as those found UE in desiccation treatment: signal transduction, cell communication, and ion transport (***Supplementary Table S2, Fig S4***). At 2.5h and 8h post-radiation, the timepoints where most genes are differentially expressed compared to the controls (***Figs 1B, 1C***), respectively 906 and 724 genes were over-expressed in the three radiation conditions (***Fig 2D***). After GO enrichment analysis of these genes found in common after IR, we measured a significant enrichment of biological processes involved in the same processes following desiccation, being protein modification and oxidation-reduction (8h post IR). A distinct significant enrichment of transcription, transport and DNA repair processes both at 2.5h and 8h post-radiation was measured after IR (***Fig 2E***).

### 4. Highly over-expressed genes following desiccation and radiation

Because half of the DE genes had no GO annotation, we used a second approach to identify the potential function of proteins encoded specifically by OE genes both after desiccation and radiation, using Pfam domains, Kegg/GO ids, and blast hits.

Selecting the timepoints with the most differential responses (more distant to the controls on the PCA; ***Fig 1B***), including 1.5h rehydration after 14 days of desiccation, 2.5h post X-ray radiation with and without desiccation and 2.5h post-desiccation and Fe-ions irradiation, a total of 258 OE genes was found in common to both rehydration after desiccation and radiation (log2fc>0.5, ***Fig 3A***; ***Supplementary Table S3***). In addition to these genes, we found 260 OE genes in all rehydration response (with or without radiation, log2fc>0.5 ***Fig 3A***), and 648 OE genes common to all radiation treatments (with or without previous desiccation, log2fc>0.5 ***Fig 3A***). Amongst those genes, we were unable to get any functional annotation (blast hit or pfam of orthology) for 58 genes in the common response (22.5%), 86 within the rehydration response (33%), and 178 within the radiation core response (27.5%). Below, we focused specifically on differentially expressed genes (log2fc>0.5) with high expression levels in the treatments (TPMs>150) to describe the key genes and processes involved in the resistances to desiccation and radiation, as presented in ***Fig 3B***.

**Fig 3.**
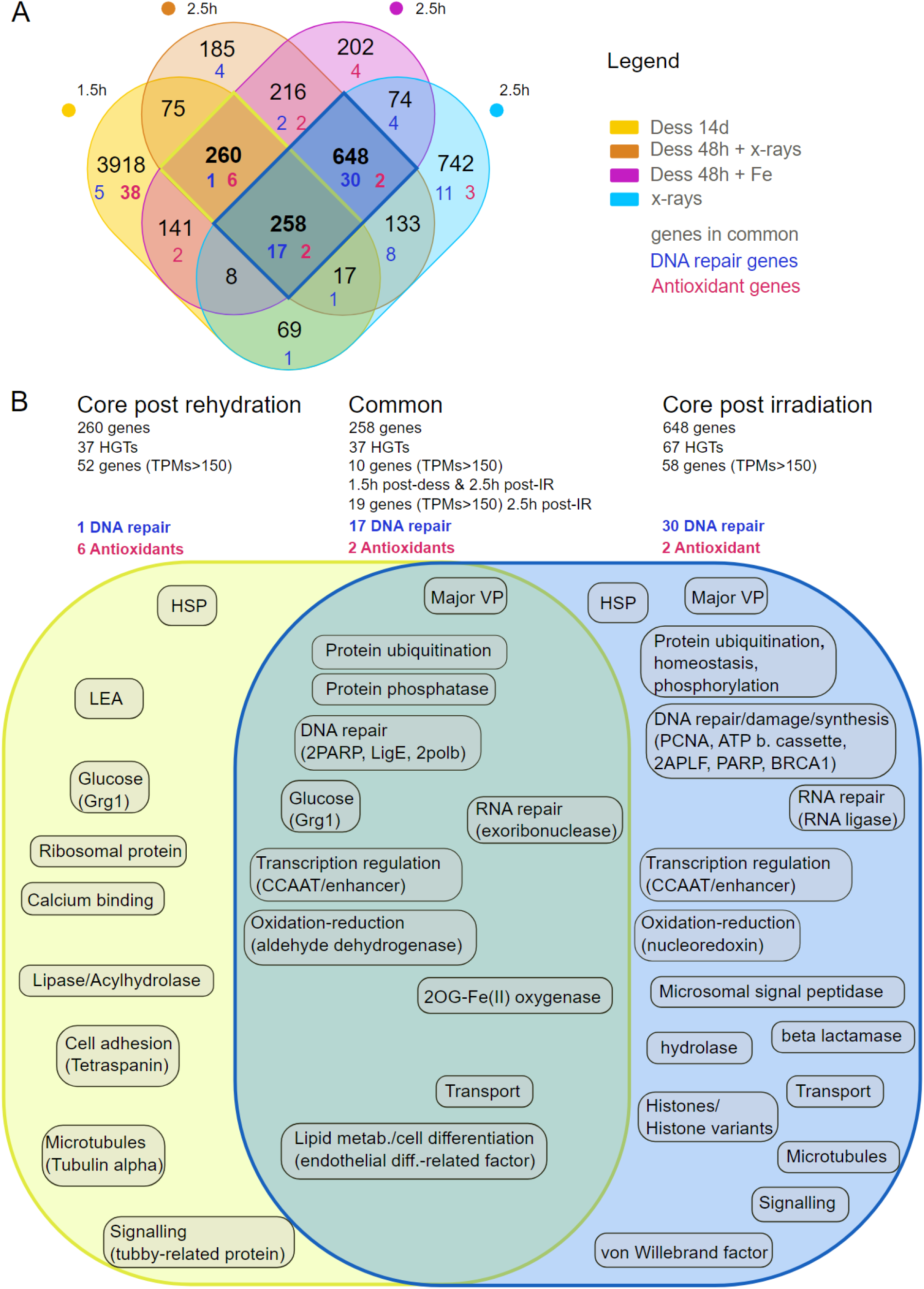
Core transcriptomic response of *A. vaga* to desiccation and radiation. A) Venn diagram of genes significantly over-expressed (OE: FDR<0.01 and log_2_foldchange>0.5 with DESeq2 and EdgeR) in *A. vaga* individuals in different studied conditions: 1.5 hours post-rehydration following 14 days of desiccation (yellow), 2.5 hours post-radiation and desiccation (Fe radiation in purple, X-ray radiation in orange), and 2.5 hours post-radiation (X-rays, blue). Genes specifically coding for DNA repair are written in dark blue and those coding for antioxidants are written in pink. B) OE genes with high level of expression (TPMs>150) and the potential function of the protein for which they code or the biological processes in which they are likely involved in the core response to rehydration (encircled in dark yellow in the Venn diagram, TPMs>150 in the condition 1.5 hours post-rehydration after 14 days of desiccation), in the core response to radiation (encircled in blue in the Venn diagram, TPMs>150 in all condition 2.5 hours post irradiation) and the genes found in both responses (common, TPMs>150 in the condition 2.5 hours post-desiccation and X-ray irradiation).

1. *Key OE genes common to both rehydration after desiccation and radiation*: we found 10 OE genes with high TPMs (>150) 2.5h post radiation (all types) and 1.5h post rehydration (without radiation) amongst the 258 OE genes of the common response to rehydration and radiation. These genes likely code for glucose-repressible gene-1 protein Grg1, and an endothelial differentiation-related factor, as well as proteins involved in ubiquitination (2 genes), transcription regulation (a CCAAT/enhancer, a transcription factor), transport, and 3 genes with unknown functions (***Fig 3B***; ***Supplementary Table S3***). The gene coding for CCAAT/enhancer likely involved in transcription regulation is the gene with the highest TPM values (TPMs>1000, 2.5h post radiation (Fe-ions or X-rays) and 1.5h post 14 days desiccation; ***Supplementary Table S3***). We decided to investigate in addition the 19 genes (log2fc>0.5, 2.5h post radiation (all types) and 1.5h post rehydration (without radiation)) also reaching TPMs>150 2.5h post radiation (all types) but not post rehydration without radiation. Indeed, desiccation is known to produce less DNA DSBs compared to IR [64]. Therefore, the potential OE genes of resistance might not reach TPMs>150 after desiccation only. The 19 additional genes were found to be likely involved in: DNA repair (2 PARP, Ligase E, 2 polymerases beta), an exoribonuclease (ERI1), an aldehyde dehydrogenase (antioxidant), a 2OG-Fe(II) oxygenase, a protein phosphatase, and one more gene involved with ubiquitination, a major vault protein, and 8 genes with unknown function.

2. *Key OE genes specific to rehydration post desiccation:* we found 52 OE genes with high TPMs (>150) amongst the 260 OE genes of the rehydration response following desiccation (with or without radiation) coding for two heat shock proteins (HSPs), LEA, Grg1, a ribosomal protein, a lipase/acylhydrolase, a protease (trypsin pfam domain PF00089), proteins with a calcium binding domain, proteins involved in signaling and microtubules, as well as 29 genes with unknown functions (***Fig 3B***; ***Supplementary Table S4***).

3. *Key OE genes specific to the radiation response:* we found 58 OE genes with high TPMs (>150) post radiation amongst the 648 genes of the radiation response, regardless of the radiation or desiccation treatment. These 58 genes are coding for proteins involved in protein modifications (ubiquitination, proteolysis, phosphorylation, homeostasis), in RNA and DNA repair/damage or synthesis, in transcription regulation (CCAAT/enhancer), in apoptosis inhibition, and in vesicle transport. We also found genes coding for HSPs, Major Vault proteins, histones and histone variants (two H2B variants, one H2A variant H2Abd2, one core histone H3, and one core histone macro-isoform X2), an RNA ligase, a nucleoredoxin, a beta lactamase, a hydrolase, a microsomal signal peptidase, von Willebrand factors and 15 genes with unknown functions (***Fig 3B***; ***Supplementary Table S5***).

Based on the 8.3% genes of non-metazoan origin (Horizontal Gene Transfer: HGTs) detected in the genome of *A. vaga* [65], we found a significant enrichment in HGTs in the different responses. For the common response, 37 HGTs amongst the 258 genes (14.3%; χ^2^=12.4, df=1, alpha=0.01; ***Fig 3B and Supplementary Table S3***) and 37 HGTs in the rehydration response following desiccation (amongst 260 genes (14.3%); χ^2^=12.01, df=1, alpha=0.01; ***Fig 3B and Supplementary Table S4***). Although we found 67 HGTs amongst the 648 genes (10.3%) in the radiation response, this did not correspond to a significant enrichment (χ^2^=3.54, df=1, alpha=0.01; ***Fig 3B and Supplementary Table S5***). Amongst these genes, 2OG-Fe(II) oxygenase, Ligase E, two RNA ligases, and genes belonging to Endonuclease/ Exonuclease phosphatase particularly retained our attention because of their high TPM values (TPMs>150). The ligase E (FUN_003353) (previously ligase K in [33] is one of the genes found in the shared response to rehydration and radiation with the highest log2foldchange (log2fc=7.32, 2.5h post-desiccation and X-ray radiation; log2fc =2.3 1.5h post 14 days of desiccation) and highest expression levels in irradiated *A. vaga* rotifers (TPMs=862, 2.5h post desiccation and X-ray radiation; TPMs=1522, 2.5h post desiccation and Fe-ions irradiation; TPMs=23, 1.5h post 15 days of desiccation) compared to the controls (1.88<TPMs<7.83) (***Supplementary Table S3***).

### 5. High constitutive expression of *A. vaga* antioxidant system enhanced by desiccation and rehydration processes

Since a significant enrichment of oxidation-reduction processes amongst OE genes in *A. vaga* individuals entering desiccation, 1.5h post-rehydration after 14 days of desiccation, and 8h post-radiation with or without desiccation was measured (***Figs 2B***, ***2E***), a functional annotation based on Pfam domains and blast search was performed for genes coding for antioxidants. In total 216 antioxidant genes were found in the entire *A. vaga* genome: 65 encoding glutathione S-transferases, 59 encoding aldo/keto reductases, 26 encoding thioredoxins, 18 encoding aldehyde dehydrogenases, 16 encoding peroxiredoxins, 10 encoding superoxide dismutases (SOD), 8 encoding isocitrate dehydrogenases, 5 encoding catalases, 5 encoding glutathione peroxidases, and 4 encoding glutathione reductases (***Supplementary Table S6***; ***Fig 4A***).

**Fig 4.**
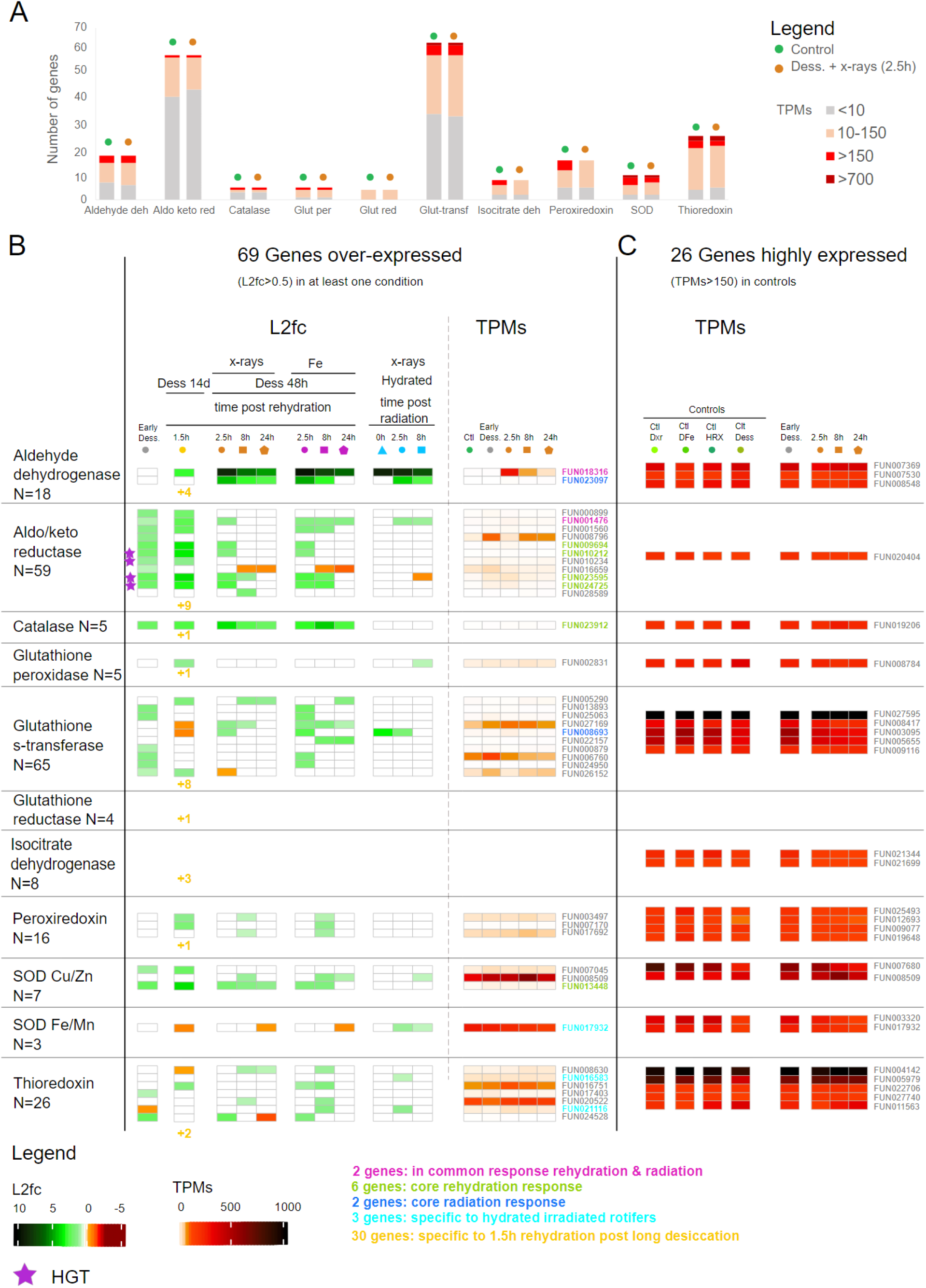
Transcriptomic response in *A. vaga* of genes coding for antioxidants. A) Number of genes coding for the different antioxidant gene families (x axis). According to the level of expression: transcripts per millions (TPMs), the number of genes is represented in different color: TPMs < 10 (gray), 10<TPMs<150 (light orange), >150 (red), >700 (dark red). The number of genes in each expression category is represented for *A. vaga* hydrated controls (green) and *A. vaga* individuals 2.5 hours post X-ray radiation and rehydration (orange). B) Heatmap of change of expression of 39 genes coding for antioxidant gene families which are at least over-expressed in one condition (log_2_foldchange > 0.5, FDR < 0.01 with both DESeq2 and EdgeR). The 30 genes found OE exclusively after 14 days of desiccation are indicated as numbers in the corresponding column. Heatmap of transcripts per millions (TPMs) for each gene is represented on the right side and with a different color code representing the level of expression. The gene IDs coding for the different kinds of antioxidants are written next to the TPMs, and Horizontal transfer genes (HGT) are represented by the violet stars. C) Heatmap representing the level of expression (TPMs) of 26 genes showing constitutive high expression (TPMs>150) in all conditions coding for antioxidants. After desiccation and radiation, the number of genes coding for antioxidants showing high TPMs does not change. The ID of the genes found in the common response to rehydration and radiation are indicated in pink (2 genes), in the common rehydration response (in green: 6 genes), in the common radiation response (in dark blue: 2 genes), specific to hydrated irradiated rotifers (in light blue). The four genes specific to desiccation and 1.5h post rehydration are indicated in yellow, same color as the 30 genes specific to 1.5h post rehydration (indicated as numbers).

We observed that many antioxidant genes were already highly expressed in the controls (TPMs>150; ***Figs 4A, 4C***). Indeed, 26 genes are highly constitutively expressed in all conditions including controls (except 2 peroxiredoxins and 1 isocitrate dehydrogenase having TPM values slightly lower than 150 post-radiation, which we considered nonetheless): three aldehyde dehydrogenases (different than the ones which are OE), one aldo/keto reductase, one catalase, one glutathione peroxidase, five glutathione s-transferases, two isocitrate dehydrogenases, four peroxiredoxins, four SOD (2 Fe/Mn, 2 Cu/Zn) with 2 of them being also OE at some timepoint post-radiation treatments, and five thioredoxins. Amongst these 26 genes, four show even higher levels of expression both in controls and desiccation/radiation treatments (TPMs>700): one superoxide dismutase (SOD Cu/Zn), two thioredoxins, one glutathione S-transferase (***Fig 4A***, ***4C, Supplementary Table S6***).

After stress, the number of those highly expressed (TPMs>150) antioxidant genes did not increase (***Figs 4A, 4B***). Still, we found 69 genes being OE in at least one condition compared to the control (***Fig 4B***). In accordance with the observed significant enrichment of oxidation-reduction processes (***Fig 2B***), the entry in desiccation and the rehydration after long periods of desiccation led to more OE antioxidant genes than radiation only (***Fig 4B***). Indeed, 1.5h rehydration post 14 days desiccated condition had the highest number of antioxidant OE genes, 48 in total (***Supplementary Table S7***), with 30 genes found to be OE exclusively in this condition (indicated as number on ***Fig 4B***). Amongst the 30 genes found OE exclusively post rehydration without radiation, we notably found two other aldehyde dehydrogenases (FUN_007530, FUN_008548) and one catalase (FUN_019206) highly expressed in the controls (respectively 150>TPMs>200; 250>TPMs>300; 190<TPMs<270 ***Fig 4A***, ***4C***) but reaching even higher TPMs 1.5h post rehydration (respectively, TPMs=303; TPMs=400, TPMs=489; ***Supplementary Table S7***). Within the remaining of the 48 genes, 4 genes were found both only in *A. vaga* individuals entering desiccation and in those being rehydrated for 1.5h after 14 days of desiccation, and 14 genes were found as well in some other condition of radiation (all time points included; ***Fig 4B***).

Only two of these genes were found in common to all the different conditions 2.5h post stress: one aldehyde dehydrogenase (already mentioned in the common response), and one aldo/keto reductase (***Supplementary results*** for more details). In the core response to rehydration we found 6 antioxidant genes (***Figs 43A, 4B***): 4 coding for aldo/keto reductases, one for a catalase, and one for a SOD (Cu/Zn). However, only 2 antioxidant genes were found in the core radiation response (2.5h post radiation): another gene coding for an aldehyde dehydrogenase and a glutathione s-transferase. A SOD (Cu/Zn) was also found in the core radiation response but 8h post-radiation (***Fig 4B***).

We found some responses specific to the treatments. For instance, less antioxidant genes were OE in hydrated conditions exposed to X-rays (7 genes OE 2.5h post X-rays) than in desiccated conditions (12 genes OE 2.5h post desiccation and X-rays). One gene, though, a SOD Fe/Mn seems to be specific to irradiated hydrated conditions as it was OE 2.5h and 8h post X-rays, even if this gene is already constitutively highly expressed in all conditions (***Fig 4B***, ***4C***). Moreover, Fe-ions radiation triggered an OE of more antioxidant genes than X-rays: 27 versus 19 genes (all time points included), with nine additional genes: 5 glutathione s-transferases, 1 peroxiredoxin, 2 thioredoxins, 1 aldo/keto reductase (***Fig 4B***).

#### Genes involved in NHEJ and BER are highly expressed post-rehydration and post-radiation

Since we found a significant GO enrichment in DNA repair biological processes in the core response to radiation, we characterized genes coding for proteins involved in DNA repair within the genome of *A. vaga*. We identified 222 DNA repair genes in *A. vaga* based on KEGG ids, Pfam domains, and genes previously identified in [33], and grouped them by repair pathway: 23 genes in NHEJ, 29 genes in HR, 46 genes in BER, 49 genes in NER, 6 genes in MR, 3 genes in alternative-NHEJ, 19 genes in DNA replication, 26 genes in cross pathways, and 21 genes with functions related to DNA repair (***Fig 5A***; ***Supplementary Table S8***).

**Fig 5.**
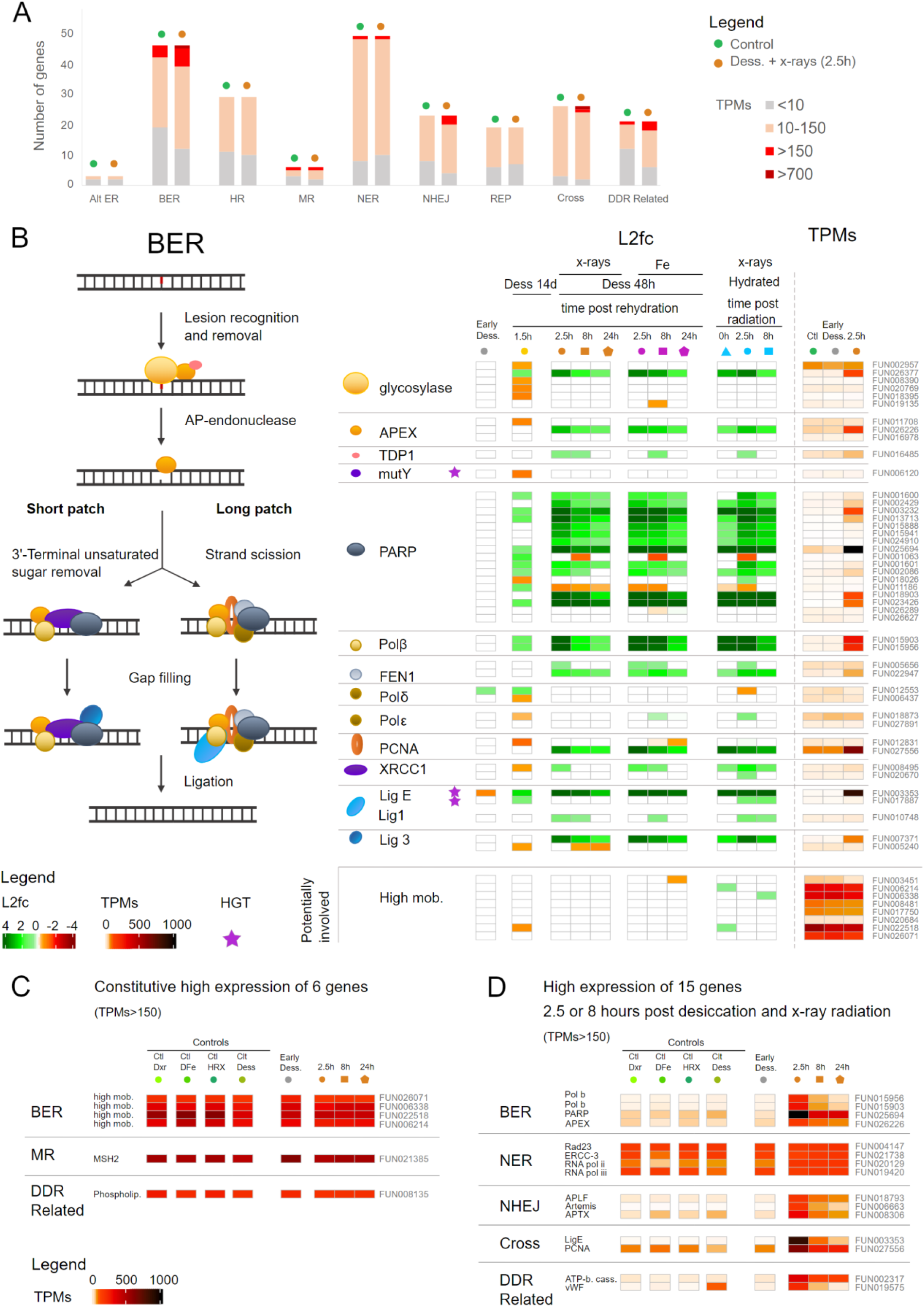
Transcriptomic response in *A. vaga* of genes involved in DNA repair pathways. A) Number of genes coding for genes involved in DNA repair pathways: Alternative Excision Repair (Alt ER), Base Excision Repair (BER), Homologous Recombination (HR), Mismatch Repair (MR), Nucleotide Excision Repair (NER), Non-Homologous End-Joining (NHEJ), DNA Replication (REP), Cross-Pathways (Cross), Damages-DNA Repair related pathways (DDR-related). According to the level of expression (transcripts per millions (TPMs)), the number of genes is represented in different colors: TPMs<10 (gray), 10<TPMs<150 (light orange), >150 (red), >700 (dark red). The number of genes in each expression category is represented for *A. vaga* hydrated controls (green) and *A. vaga* individuals 2.5 hours post X-ray radiation and rehydration (orange). B) BER pathway showing numerous genes being over-expressed (L2fc > 0.5) and some with high TPMS after desiccation and/or radiation. This figure represents the different actors of the BER pathway and their interaction, actors being represented by different symbols. The heatmap of the log_2_foldchange shows of the level of expression of BER genes change compared to the expression level in the controls. The color is not white when the gene is significantly (FDR < 0.01 with both DESeq2 and EdgeR) over- (log_2_foldchange>0.5 in green) or under-expressed (log_2_foldchange<- 0.5 in orange or red). Genes potentially involved in this pathway are indicated in the bottom of the figure. The investigated conditions are: individuals entering desiccation (gray), after 14 days desiccation and 1.5 hours rehydration (yellow), after desiccation and X-rays radiation (orange), after desiccation and Fe radiation (purple), after X-ray radiation without desiccation (blue). The investigated time points are represented as follow: 0 hours (triangles), 1.5 or 2.5 hours (circles), 8 hours (squares), 24 hours (pentagons). Heatmap of transcripts per millions (TPMs) for each gene is represented on the right side and with a different color code representing the level of expression. The gene IDs coding for the different kinds of BER genes are written next to the TPMs, and Horizontal transfer genes (HGT) are represented by the violet stars. C) Heatmap representing the level of expression (TPMs) of 6 genes coding for proteins involved in DNA repair pathways (BER, MR, DDR-related) showing constitutive high expression (TPMs>150) in all conditions. D) Heatmap representing the level of expression (TPMs) of 15 additional genes coding for proteins involved in DNA repair which have a high gene expression level after radiation and desiccation.

Unlike numerous genes encoding antioxidants that are constitutively expressed in the controls, only six genes involved in DNA repair have a high expression level (TPMs>150) in all controls: four genes coding for high mobility group proteins potentially involved in BER [66], MSH2 involved in MR, and a phospholipase involved in related DNA repair processes (***Figs 5A; 5C***).

We found 17 DNA repair genes in the common response to rehydration and radiation and 30 additional genes in the core response to radiation (***Fig 3A*** (genes indicated in blue); ***Figs 5B***, ***Fig 6***, ***Figs S5***, ***S6***, ***S7***, ***S8, Supplementary Table S8***). Most genes being OE 2.5h post-radiation belong to BER (21/53) and NHEJ (12/23) pathways, unlike genes from HR (6/40), NER (8/63), or MR (4/31), taking into account genes involved in cross-pathways (***Figs 5A, 5B, 6***, ***Figs S5***, ***S6***, ***S7***, ***S8***). The few genes with high log2foldchange values in HR, NER and MR are those coding for proteins involved in cross-pathways (e.g., PCNA; Ligase E), whereas genes coding for proteins exclusively involved in BER (e.g., Polymerases β, PARPs) and NHEJ (e.g., KU80; Artemis; DNA-PKcs; APLF; APTX) reach high log2foldchange values (***Figs 5B, 6, Figs S5***, ***S6***, ***S7***, ***S8***). Most of these genes also reach high TPMs (>150), in particular one gene coding for PARP reached TPM values >1000 2.5h post-radiation. We noticed that some actors in BER (e.g., glycosylase, APEX, PCNA and Ligase 3) and in NHEJ (e.g., KU70, KU80, Artemis, Polymerase λ) had only one copy over-expressed after radiation, whereas other actors had multiple copies over-expressed (e.g., XRCC4, APLF, PARP, Polymerase β, FEN1) (***Figs 5B, 6***).

**Fig 6.**
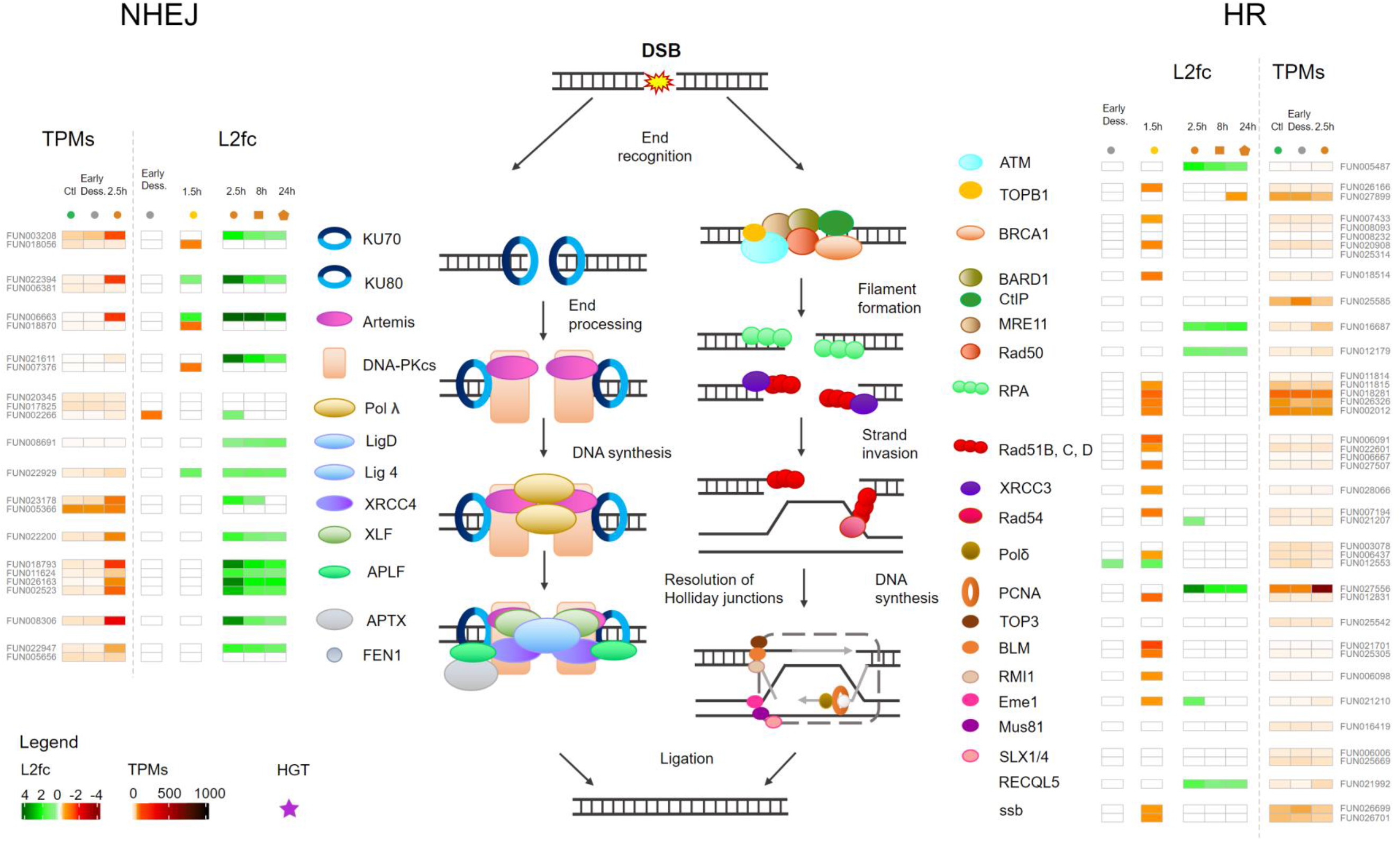
Expression of genes involved in the Non-Homologous End-Joining (NHEJ) and Homologous Recombination (HR) pathways. The figure represents the different actors, being represented by different symbols, acting in the two DNA repair pathways which repair double strand breaks (DSBs). The heatmap of the log_2_foldchange shows the change level of expression of genes coding for these actors in the different conditions in *A. vaga* individuals compared to the gene expression level in the controls. The color is not white when the gene is significantly (FDR < 0.01 with both DESeq2 and EdgeR) over- (log_2_foldchange>0.5 in green) or under-expressed (log_2_foldchange<-0.5 in orange or red). In this figure are represented only the following conditions: individuals entering desiccation (gray), after 14 days desiccation and 1.5 hours rehydration (yellow), after 2.5-, 8- and 24-hours post desiccation and X-rays radiation (orange), since the response was similar after desiccation and Fe radiation and after X-ray radiation without desiccation (all conditions represented in ***Supplementary Figures S4 and S5***). Heatmap of transcripts per millions (TPMs) for each gene is represented on the right side and with a different color code representing the level of expression. The gene IDs coding for the different DNA repair genes are written next to the TPMs, and Horizontal transfer genes (HGT) are represented by the violet stars. This figure illustrates that DSB are likely repaired in *A. vaga* individuals by NHEJ rather than by HR.

Most of the genes found OE post rehydration without radiation were found OE post radiation, such as KU80, Artemis, Lig4, glycosylase, PARP, polβ, Ligase E (***Figs 5B, 6***) indicating similar DNA repair mechanisms for both stresses. Moreover, many genes involved in DNA repair which were OE post-radiation were slightly OE post-rehydration without radiation but were below the established thresholds for log2foldchange and FDR.

We observed for the different types of radiation the same genes involved in DNA repair being OE, with similar log2foldchange values and with, for most of them, the highest values 2.5h post-radiation (***Figs 5B***, ***6***, ***Figs S5***, ***S6***, ***S7***, ***S8***). Therefore, we represented TPMs only 2.5h post-desiccation and X-ray radiation in ***Figs 5A***, ***5B***, ***5C***, and ***5D***. The only difference we noticed was that some genes OE post X-rays radiation were significantly OE 8h post Fe-ions irradiation, and not 2.5h (e.g., TDP1, Ligase 1 in BER, ***Fig 5B***; Ligase 4, Ligase D in NHEJ, ***Fig S5***; XPC and CSB in NER, ***Fig S7***).

## Discussion

Transcriptomic analyses of the bdelloid rotifer *A. vaga* facing desiccation and IR allowed us to determine which genes and biological processes are likely involved in response to these stresses. Moreover, this allowed identifying genes commonly responsive to both stresses and some candidates that appeared specific to a peculiar response (summarized on ***Fig 7***). We mainly discuss below enriched biological processes, and overrepresented gene families with OE genes (log2fc>0.5, FDR<0.01) reaching high expression levels (TPMs>150) after IR or desiccation.

**Fig 7.**
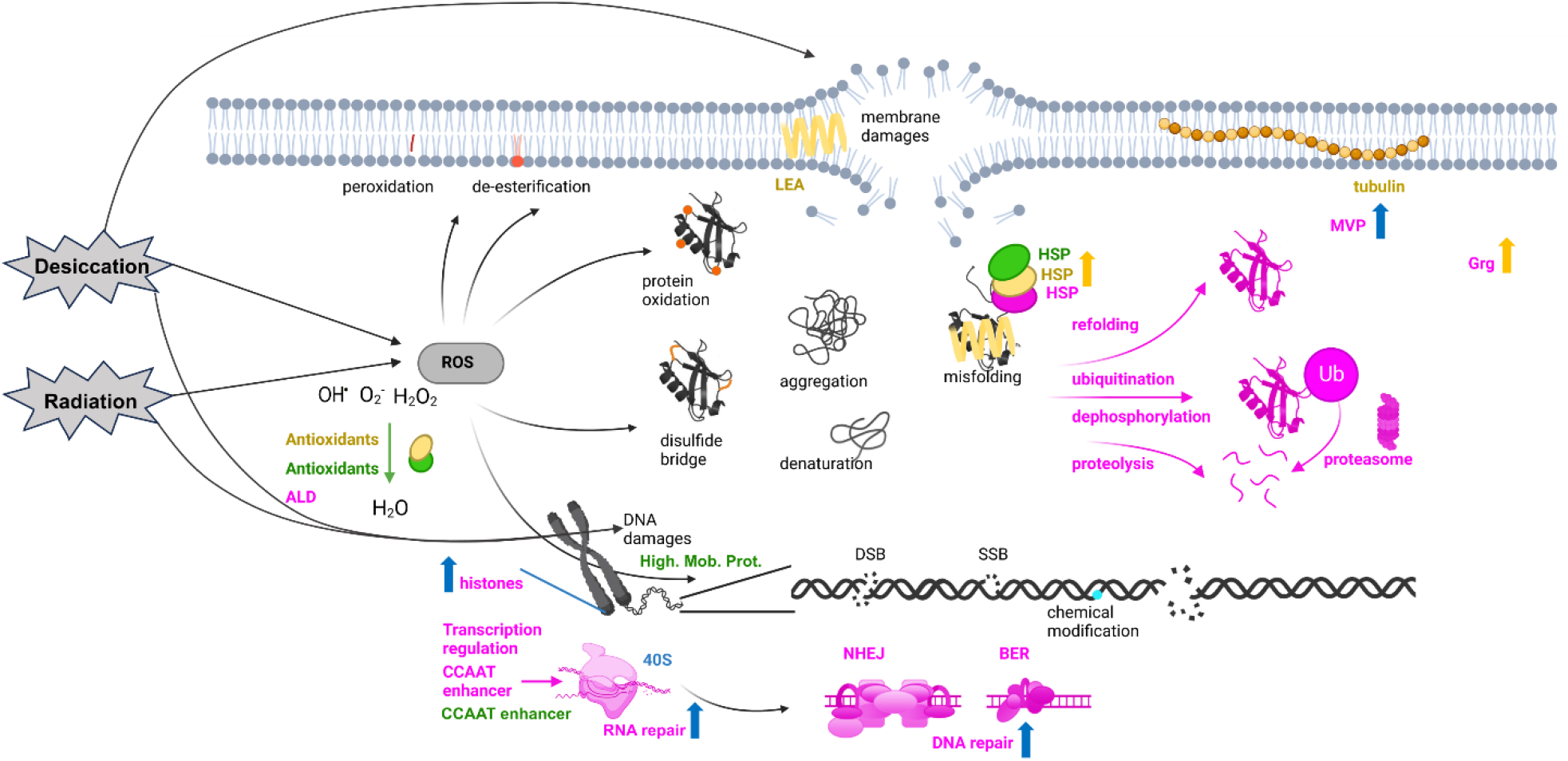
Summary of common and specific responses to desiccation and ionizing radiation. Common damages known to be induced by desiccation and ionizing radiation: membrane damages, ROS production leading to phospholipid peroxidation and de-esterification, protein damages (oxidation, disulfide bridge, denaturation, aggregation, misfolding), and DNA damages (Double Strand Break: DSB, Single Strand Break: SSB, chemical modifications). The major molecular responses observed in *Adineta vaga* post-rehydration (with or without radiation; in yellow) and post-radiation (in blue), or common to both (in pink). Genes coding for some antioxidants, HSPs, high mobility proteins, and one CCAAT enhancer were found to be constitutively highly expressed (in green) in all conditions including controls. We found genes coding for HSPs, Major Vault proteins (MVP), Glucose repressive proteins (Grg1), aldehyde dehydrogenase, regulation factors such as CCAAT enhancer, histone variants genes involved in protein modifications like ubiquitination, in proteolysis, DNA repair (mostly NHEJ and BER pathways) and RNA repair, being OE in the common response to rehydration and irradiation. Specific to rehydration, we found OE genes coding for Late Embryogenesis Abundant proteins (LEA), Heat Shock Proteins (HSPs), tubulin, Glucose repressive proteins (Grg1), and more antioxidants. Radiation increases the number of OE genes coding for histone variants, RNA/DNA repair proteins, and MVPs.

### 1. Common transcriptomic response of *A. vaga* to rehydration and radiation

Both desiccation and radiation lead to DNA/protein damages and ROS formation [9–12]. These damages and ROS can lead to compromised cellular function and even cell death, emphasizing the need for a swift and rapid expression of protection and repair mechanisms.

Similar damages, including DNA DSBs [12, 19], caused both by desiccation and IR likely explain why we found a common transcriptomic response in the bdelloid rotifer *A. vaga* (***Fig 7*** in pink) both post-rehydration (1.5h) and radiation (2.5h) with the oxidoreduction and protein modification processes being enriched (***Figs 2B*** and ***2E***) and 258 genes being commonly OE including 17 genes involved in DNA repair (***Figs 3A*** and ***3B***). For most of the OE genes, the over-expression was at the highest early after stress exposure (timepoints 1.5h or 2.5h post rehydration or radiation), indicating a fast transcriptomic response from the bdelloid rotifer *A. vaga* upon exposure, probably contributing to its high resistance and resilience.

At protein level, cells can scavenge ROS with antioxidants to prevent oxidative damage in general [67], or protect proteins with chaperones like HSPs [68, 69]. In general, molecular chaperones play a crucial role in protein homeostasis (proteostasis) by regulating protein quality control, folding and turnover, preventing newly synthesized proteins to aggregate into non-functional structures [68, 70]. While antioxidant genes appear constitutively expressed in *A. vaga* (***Figs 4A, 4C, Fig 7*** in green), two genes coding for antioxidants, one aldehyde dehydrogenase and one aldo/keto reductase were found highly OE in the common response (***Fig 5B***), suggesting the presence of a general mechanism of oxidative stress prevention. Moreover, we found evidence of several HSP (e.g., HSP70) coding gene constitutively highly expressed (TPMs>150; ***Fig 7*** in green; ***Supplementary Table S1***) and some genes over-expressed in the common response (***Fig 3B***). In general however, more genes coding for HSPs and antioxidants were identified in the rehydration post-desiccation response when compared to radiation only (***Fig 4B***).

When antioxidant and chaperones systems fail to respectively protect the proteins or to refold a denatured protein, proteins can be modified. In our transcriptomic analysis on *A. vaga* we also found genes and enriched biological processes OE after both stresses related to protein modifications and ubiquitination (***Figs 2E***; ***Fig 3B***). These protein modifications can cause change of a protein activity or function (e.g., dephosphorylation) or mark targeted proteins for degradation [71]. Ubiquitination often marks misfolded proteins for proteolysis involving the 26S proteasome, a large multiprotein complex [71–73]. Proteins involved in protein ubiquitination were also found OE in other desiccation resistant species in response to rehydration (e.g., in tardigrade: [74]). Interestingly, ubiquitin-dependent signaling processes have also been shown to play a role in the choice of the double strand break (DSB) repair pathway (HR or NHEJ) [75] and in the re-establishment of genome integrity after DSBs [76].

While proteins appear to be protected from oxidative damage in *A. vaga* during radiation (see also [30]), DNA is not protected and damages accumulate with prolonged desiccation and higher doses of radiation [12]. DNA double strand breaks (DSBs), but also single strand break (SSBs), and base damages are known to be caused by direct or indirect damages following radiation [57, 58, 77, 78] and were indeed observed in *A. vaga* [12, 19]. DNA DSBs were also detected during prolonged desiccation in *A. vaga* but less DSBs were visible on pulsed-field gel electrophoresis (PFGE) than after 500 Gy X-ray radiation [12, 19]. High radiation doses were chosen in this study to detect all actors involved in radiation resistance. Here, we found 17 genes to be involved with DNA repair in the common transcriptomic response (***Fig 3A***). The high radiation levels (500 Gy) inducing more DSBs than desiccation [19] probably explains why we found more repair genes OE in the core response to radiation (30 additional genes; ***Figs 3A***; ***Fig 7*** in blue) and why one of the most enriched biological processes post-radiation is DNA repair (***Fig 2E***). Many of these 30 additional repair genes were slightly OE post-rehydration without radiation, but with log2fc and FDR values close to the chosen thresholds (***Supplementary Table S8***).

Unlike antioxidants for which many genes are already highly expressed in the controls, we found only six genes involved in DNA repair which are highly constitutively expressed (TPMs>150 in all conditions, including controls), notably four high mobility proteins (***Fig 5C***; ***Fig 7*** in green). High mobility proteins have a role in the chromatin structure, transcriptional regulation but also seem to enhance DNA repair [79]. Indeed, the absence of high mobility group protein B1 (HMGB1) was shown to increase mutagenesis, decrease cell survival, and alter chromatin reorganization after DNA damage [79]. It was also shown that HMGB1 was involved with telomere homeostasis and prevented DNA damage induced by radiation in human breast cancer cells [80]. Like antioxidants being highly expressed in the control conditions, the constitutive expression of genes coding for high mobility proteins could be advantageous in *A. vaga* for chromatin reorganization, transcriptional regulation and improved DNA repair during the numerous unpredicted episodes of desiccation in their semi-terrestrial environments.

We found that the DNA repair proteins encoded by genes OE post desiccation and/or radiation were mostly involved in BER and NHEJ DNA repair pathways. Some actors seem particularly important (as they reached high expression levels) such as PARP and two polymerases beta in BER pathway (***Fig 5B***), APLF and APTX in NHEJ pathway (***Fig 6***), and DNA ligase E and PCNA involved in cross pathways (***Figs 5B, 6***). Specifically, the PARP gene family may play a key role for the bdelloid rotifer resistance as we found a high number of over-expressed copies: 17 genes are OE 2.5h post-radiation (high and low LET IR) and 9 genes are OE post-rehydration after 14 days of desiccation (***Fig 5B***). PARPs are involved in BER pathway, notably for recruiting other proteins of this pathway [81]. It has already been shown that PARPs are important to deal with radiation DNA damages as their expression level increases in carcinogen-treated cells or in cells exposed to radiation or DNA damaging agents [82–84]. Besides DNA repair, PARPs appear to be involved in other cellular processes such as the modulation of chromatin structure, transcription, replication, recombination, regulation of membrane structures, cell viability, cell division, and actin cytoskeleton [85], explaining perhaps their over-expression.

Hecox Lea and Mark Welch [33] already highlighted that genes of the DNA repair pathways are upregulated in *A. vaga* individuals desiccated for 7 days and rehydrated for 1h, with repair actors of different pathways involved. This included ligase E (named ligase K in [33]) involved in DNA ligation during DNA repair (also found OE in this study) but also genes coding for APLF (found in this study but 0.01<FDR<0.05, 1.5h post-rehydration) and polymerase lambda of the NHEJ pathway (also found OE in this study). However, they also found BLM (HR pathway) to be OE (not the case in our study), whereas we found in addition KU80 or Artemis from the NHEJ pathway (***Fig 6***, ***Fig S5***) and several actors of the BER pathway including a glycosylase, 9 PARPs, one polymerase delta (***Fig 5B***) to be OE 1.5h post-rehydration. Since IR or prolonged desiccation (14 vs. 7 days) causes more DNA damages, we were able in our study to amplify the response and found that the major DNA repair pathways in *A. vaga* are BER and NHEJ. Moreover, the log2fc values might have been bigger if we measured the gene expression levels of rehydrated rotifers at other time points (e.g., 2.5h, and 8h).

The genes found OE in bdelloid rotifers in this study associated with specific DNA repair pathways were different from the repair genes reported in other radiation and desiccation resistant species. In the bacterium *D. radiodurans* proteins involved in NER and HR were upregulated post-desiccation and post-radiation [13]. In *P. vanderplanki* insect larvae proteins of HR (Rad23 and Rad51) were upregulated both after desiccation and high- and low-LET radiation [86], while in tardigrades all major repair pathways appear expressed [87]. This suggests that distinct DNA repair pathways are active in different desiccation resistant species.

DNA DSB repair is known to require, in many eukaryotes, the conserved histone variant of canonical H2A, H2AX, which is absent from studied bdelloid rotifer species [88]. In response to DSBs, H2AX becomes phosphorylated in its C-terminus, triggering the retention and accumulation of repair and checkpoint proteins to the DNA breaks [88]. Amongst other H2A variants in the *A. vaga* genome, some were OE after both stresses (e.g., H2Abd1; TPMs>100 2.5h post radiation; ***Supplementary Table S3***; ***Fig 7***). In addition to H2Abd1, we found an increased expression of other histones following radiation, such as two H2B variants, the other identified H2A variant (H2Abd2), one core histone H3, and one core histone macro-isoform X2 (TPMs>200, 2.5h post-radiation; ***Supplementary Tables S3 and S4***). We do not know, so far, the exact role of these histone variants in the resistance of *A. vaga*. In general, core histones and its variants play a role in chromatin (de)compaction and in the maintenance, replication and expression of the genome, which is important when numerous genes are transcribed in response to desiccation and/or radiation stress (with over 1000 genes over-expressed post-stress). We therefore also identified actors involved in DNA replication, RNA synthesis, and RNA maturation processes: 40S ribosomal protein and nucleolin-like isoform protein found OE and highly expressed in the core response to radiation (***Fig 3B, Fig 7*** in blue).

Besides DNA damages, RNA damages likely also occur during desiccation and radiation but are so far less studied [89]. However, RNA might be even more prone to modifications and degradation compared to DNA notably because of their single stranded structure [90], the non-interaction with proteins or nucleosomes protecting the DNA [91], and their high abundance in the cell (80–90 % of the total nucleic acid composition; [90]). While less lethal than genome mutations, their non-repair could result in delayed or faulty translation and inactive proteins, or dysregulation of gene expression [92]. Indeed, oxidative RNA damage has been associated with neurodegenerative disorders [92]. Here, we detected the OE of genes coding for proteins involved in RNA repair, RNA silencing and degradation of RNA substrates in the common response (***Fig 7*** in pink) such as RNA ligases [93] or ERI1 exoribonuclease [94, 95].

We identified OE genes belonging to distinct gene families with high log2foldchange and/or TPM values compared to the controls, suggesting that they may play a role in *A. vaga* resistance and warrant further investigation. Among those genes, we noticed the 2OG-Fe(II) oxygenase superfamily, with 49 annotated genes in the *A. vaga* genome, six over-expressed in the common response to rehydration and radiation (***Supplementary Table S3)*** and seven others over-expressed under all radiation conditions (***Supplementary Table S5***). This gene family is known to be involved in various processes essential for stress resistance, such as protein modification, DNA and mRNA repair, and synthesis of secondary metabolites [96, 97]. Second, we found the major vault proteins (MVPs) with five OE genes: one found in the common response to rehydration and radiation (TPMs>150, 2.5h post radiation), and four in the common response to the different radiation types (two with TPMs>150, 2.5h post radiation). Such proteins have been linked to stress response in other studies, such as drug resistance, DNA repair [98] and response to DNA-damaging agents like IR [99]. Finally, two OE genes in the shared response and reaching high TPMs coded for proteins characterized by the domain “bZIP transcription factor” (PF00170). This domain is found in proteins regulating different physiological processes (e.g., adipogenesis, response to oxidative stress, in surveillance immunity, or cancer progression; [100]). One of these genes, likely coding for a CCAAT/enhancer-binding protein, reached the highest TPM values in the common response (TPMs>1000, 2.5h post radiation or 1.5h post-rehydration) (***Supplementary Table S3***). We hypothesize that this putative transcription factor likely triggers the downstream upregulation of several genes involved in the response to desiccation and/or radiation in *A. vaga*.

Finally, unlike the radio-sensitive monogonont rotifer *Brachionus koreanus* in which the p53 gene, trigger of apoptosis in mammalian cells [101, 102], is OE post gamma radiation (150 and 200 Gy; [103]), p53 gene is not present in the *A. vaga* genome. Similarly, this gene seems also absent in the cryptobiont heterotardigrade *Echiniscoides* cf. *sigismundi* [104]. The absence of p53 and apoptosis following extensive DNA damage in *A. vaga,* suggests that apoptosis is unlikely to occur in bdelloid rotifers exposed to radiation or desiccation. Instead of inducing programmed cell death when damages accumulate, regulated by p53, bdelloid rotifers seem to possess an arsenal of genes to prevent and repair the damages incurred, at least in their somatic cells that are not engaged in mitosis (bdelloid rotifers being eutelic).

### 2. Genes specific to desiccation and/or rehydration process

Although numerous genes were identified in the common transcriptomic response to rehydration following desiccation and radiation, some genes appeared to be specific to desiccation stress (***Fig 7*** in yellow). Desiccation not only induces damages and generates reactive oxygen species (ROS) similar to ionizing radiation (IR) (as shown in the tardigrade *Paramacrobiotus spatialis*, [105]), it also affects the physiological state of the organism (hydrated vs. desiccated). Here, we showed that the dehydration process leading to the complete desiccation of *A. vaga* was characterized by a decrease of metabolic activity (many under-expressed genes) with significant enrichment amongst UE genes of signal transduction, transport, and cell communication processes **(*Fig 2A*)**. Similar patterns of gene under-expression were also observed during anhydrobiosis in the limno-terrestrial tardigrade *Milnesium tardigradum* [74].

Despite a general trend of reduced expression during dehydration (***Fig 1C***), desiccation survival requires complex adaptations at morphological, physiological, and molecular levels to protect cellular components from desiccation-induced damage or facilitate their repair upon rehydration and therefore many genes were found OE post-rehydration following 14 days desiccation (***Fig 1C***). Some proteins seem particularly important in desiccation entry and the rehydration process following desiccation as we found 52 OE genes with high expression levels (TPMs>150) specific to post rehydration (***Fig 3B***). These OE genes code for cell-adhesion proteins and microtubules, chaperones such as Late Embryogenesis Abundant (LEA) proteins and Heat Shock Proteins (HSPs), Glucose-repressible proteins, lipase and proteins involved in signaling, or characterized by a calcium binding domain (***Fig 3B***), in addition to what is already described for the shared response. We describe below the genes for which we can interpretate their potential functions regarding desiccation resistance.

Genes coding for cell-adhesion proteins and microtubules (e.g., tubulin) that were over-expressed during desiccation entry and post-rehydration likely play a role in cell shape reorganization during water loss (desiccation) and rehydration (***Fig 7*** in yellow). Previous studies have reported alterations in the cytoskeleton and cell adhesion complexes in desiccated and rehydrated bdelloid rotifers [46]. Similar genes were also found to be over-expressed after rehydration in the tardigrade *Milnesium tardigradum* [74]. In agreement with these findings, multiple homologs of such genes were reported in the genome of the bdelloid rotifer *A. ricciae*, supporting their importance in desiccation-resistant bdelloids [106].

Several anhydrobiotic species produce disaccharides, such as trehalose, which are known to replace water and vitrify the cytoplasm [47, 107–112]. Surprisingly, trehalose has never been detected in bdelloid rotifers. In our study, we confirmed the over-expression and high abundance of trehalase genes (involved in the trehalose metabolic pathway, enriched during desiccation entry) potentially correlated with the absence or low abundance of trehalose in desiccated bdelloids [32, 48]. However, we revealed that carbohydrate metabolic processes were enriched during the entry in desiccation (and only then) in bdelloid rotifers (***Fig 2B***). Genes involved in carbohydrate metabolic processes could play a role in the production of carbohydrates instead of trehalose to vitrify the cytoplasm, stabilizing the membranes and macromolecules upon desiccation [113–115]. In addition, the over-expression of such genes in bdelloid rotifers entering desiccation may mediate their metabolism during desiccation-induced shutdown because carbohydrates serve as a major source of energy and carbon. The expansion of carbohydrate-related genes in *A. vaga* and *A.ricciae* genome has been attributed to adaptations to diverse food sources, which in turn broadens their ecological niches, as observed in human gut microorganisms [12, 31, 43].

We found that genes coding for chaperones are highly OE following desiccation (***Fig 7*** in yellow). First, hydrophobic LEA proteins are highly expressed and likely play a similar role as trehalose during desiccation and rehydration processes in bdelloid rotifers. These proteins were already detected to be upregulated in the bdelloid rotifer *Philodina roseola* during desiccation [49], in desiccation resistant plants [116], in bacteria [13, 117], and in other desiccation-resistant metazoans [14, 52, 118–122]. LEA proteins likely stabilize the cellular membranes in the absence of water (as suggested by [49, 50, 52, 123, 124], protect against protein aggregation [125], and have similar functions as non-reductive sugars in vitrification of the cytoplasm [52, 126, 127]. LEA proteins have also been shown to protect crucial metabolic enzymes and were suggested to act as chaperone-like [128–131]. Second, other types of chaperones highly expressed during the rehydration process of *A. vaga* (not OE during desiccation entry) were the heat shock proteins (HSPs), a few of them also found in the common response to radiation and rehydration. Some of the OE HSPs already reached high expression levels in control condition (TPMs>300, ***Supplementary Table S4***; e.g., FUN_27450 coding for a HSP70, TPMs= 2239), but even higher expression levels post-rehydration (e.g., TPMs= 3350 1.5h post rehydration following 14 days desiccation). Radiation also triggered the over-expression of genes coding for HSPs but not as high as desiccation. HSPs are known to act as protein chaperones and therefore likely help nascent and misfolded proteins to gain their correct conformation [69–71]. The high number of HSP genes (74 according to Pfam domain annotations) detected in *A. vaga* genome, their differential expression upon desiccation/rehydration processes and the high level of transcripts reported in this study, may support a central role of HSPs in the desiccation and radiation resistance. Indeed, such proteins were found to be over-expressed in other desiccation-resistant species to deal with such stresses [51, 132–135].

Upon rehydration, one gene family particularly retained our attention because several copies were over-expressed and reached high levels of expression (TPMs>150). These were annotated with the Pfam domain Glucose-repressible protein Grg1 (PF11034). Amongst the eight genes annotated with this Pfam domain Grg1 (PF11034) in *A. vaga* genome, six of them were OE with high TPMs (TPMS>150) post-rehydration (five also at desiccation entry), one being found in the common response to rehydration and radiation (***Fig 7***). One gene was not OE but showed high constitutive TPMs in the controls (***Fig 7*** in green; ***Supplementary Table S3***). This gene family was already found over-expressed in the bdelloid rotifer *A. ricciae* entering desiccation [43], but was never reported in other anhydrobiotic metazoans, including tardigrades, nor in the desiccation-resistant bacterium *D. radiodurans*. Although these genes were not predicted as acquired via HGTs in *A. vaga* using the Alienomics pipeline [34, 65], the best blast matches were among fungi (e.g., *Microbotryum lychnidis-dioicae*, *Cladophialophora carrionii* with e-value between 1e-12 and 5e-4) which was concordant with the gene tree inferred from [43]. It is interesting to note that genes coding for Grg1 are expressed in fungi during the conidiation process when there is nutrient deprivation and desiccation [136]. We hypothesize that these Grg1 genes could have been horizontally acquired from fungi and retained for their plausible role in desiccation resistance (and glucose absence) characterized by a metabolic arrest in bdelloid rotifers.

### 3. Antioxidant role in desiccation and radiation resistances

Both desiccation and IR result in ROS production [30, 35–37] that induces damages if not mitigated by antioxidants. Our dataset reveals that major antioxidant genes are highly expressed in both control and desiccated/irradiated samples (***Fig 7*** in green), with 26 genes showing high expression levels (TPMs>150; ***Fig 4A, 4C***). This suggests that the constitutive expression of these antioxidants in *A. vaga* individuals may help them cope with oxidative stress, as they often encounter unpredictable desiccation events in their semi-terrestrial environments. Constitutive expression of protective proteins was also suggested in tardigrades [137].

We however observed that there is a higher OE of antioxidant genes following desiccation (48 OE genes; yellow condition in ***Fig 4B***; ***Fig 7*** in yellow) than radiation only (9 OE genes when combining all time points; blue condition in ***Fig 4B***). This link between anhydrobiosis and the antioxidant response was already emphasized by the many gene copy numbers of antioxidants found in the genome of *A. vaga* [31]. This observation suggests either that the ROS concentration increases following desiccation or that the number of antioxidants decreased during desiccation and were re-expressed following rehydration.

Within the antioxidants specific to the core rehydration response (indicated in green in ***Fig 4B***), we found four aldo/keto reductases, one catalase, and one Copper/Zinc superoxide dismutase (SOD). While this Copper/Zinc SOD (FUN_013448) and another one (FUN_007045) were found OE post desiccation and after long rehydration, an Iron/Manganese SOD (FUN_017932, based on blast hit, likely a Manganese SOD) was found to be OE only in hydrated irradiated rotifers (***Fig 4B***), even if it is already highly constitutively expressed (***Fig 4C***). This observation suggests an important role of that SOD for the resistance of *A. vaga* in desiccated and even more in hydrated conditions when irradiated. An over-expression of genes coding for catalase, glutathione reductase, glutathione peroxidase, and SOD have also been reported for tardigrades during desiccation [105, 138, 139].

Two genes however seem to be key actors in *A. vaga* to deal with ROS and oxidative stress following radiation as they are highly OE post-radiation: one aldehyde dehydrogenase found in the common response to rehydration and radiation, and a nucleoredoxin found in the core response to radiation (not considered in this study and in [31] as an antioxidant gene family). These genes, reaching higher expression level post-radiation, might be involved in additional processes other than their protective role against ROS. Indeed, the nucleoredoxin (TPMs>1000, 2.5h post radiation) might be required to keep the integrity of the antioxidant system, as it has been shown in plants [140]. It was also shown in the tomato that nucleoredoxin can act as a positive regulator increasing antioxidant capacity and inducing HSPs to protect cells against oxidative damage and protein denaturation during heat stress [141]. The aldehyde dehydrogenase seems to be important for multiple organisms in response to different stresses as it has been observed to be upregulated in bacteria (environmental and chemical stress), plants (dehydration, salinity, and oxidative stress), yeast (ethanol exposure and oxidative stress), *Caenorhabditis elegans* (lipid peroxidation), and mammals (oxidative stress and lipid peroxidation) [142]. Its higher expression post-radiation (TPMs>200) might be due to its potential role in the activation of DSBs resistance and DNA repair [143, 144]. This could explain why this gene was highly over-expressed post-radiation compared to other antioxidant genes in the bdelloid rotifer *A. vaga*, as X-rays lead to more DSBs than desiccation [12, 19]. In addition, we found four other aldehyde dehydrogenases (***Fig 4B***) which were over-expressed only post rehydration after long desiccation. These genes might be involved more specifically in damages caused by desiccation only.

### 4. HGTs have contributed to the evolution of resistance mechanisms in bdelloid rotifers

Genomes of bdelloid rotifers such as *A. vaga* were described to contain an unusually high proportion of HGTs [10, 12, 31, 34, 42, 145], determined to be 8.3% of protein-coding genes [34]. Frequent desiccation events might explain this high percentage because membranes are leakier and DNA fragmented, facilitating horizontal gene transfer events. It was previously hypothesized that HGTs in bdelloid rotifers may play a role in their extreme resistance to various stress [43, 145–147], unlike other species resistant to desiccation and/or radiation (e.g., [42, 104, 148]. However, this hypothesis needs to be further evaluated as a high proportion of HGTs was also found in the non-desiccating *Rotaria* rotifers *(R. macrura*, and *R. magnacalcarata*; [42]). We hypothesize that some HGTs important for resistance to extreme stresses were acquired in the ancestor of bdelloid rotifers (because all tested bdelloid species have >6% of HGTs) and have provided a selective advantage, being retained. Indeed, if anhydrobiosis was the ancestral state, it explains the high proportion of HGTs acquired prior to the loss of anhydrobiosis in some non-desiccating rotifers.

In this study, we found a significant enrichment of HGTs in the common transcriptomic response to rehydration following desiccation and radiation, and in the response to rehydration reaching in both cases ca. 14% of the OE genes (***Fig 3B***). Similar results (14.2%) have been found in response to rehydration by [42]. HGTs involved with desiccation resistance have also been described previously in rotifers [33, 43, 146]. HGTs detected by the pipeline Alienomics [34], such as 2OG-Fe(II) oxygenases, Ligase E, and two RNA ligases, particularly retained our attention because of their high TPM values and their potential role as key actors to resist desiccation and radiation. For instance, the ligase (FUN_003353) characterized as ligase E by ***Nicolas et al. (in press)*** (previously ligase K in [33]) is one of the genes found in the shared response to rehydration and radiation with the highest log2foldchange (log2fc=7.32, 2.5h post desiccation and X-ray radiation; log2fc =2.3 1.5h post 15 days of desiccation) and highest expression levels in irradiated rotifers (TPMs>800 2.5h post-radiation; ***Supplementary Table S3***). This gene, responsible for DNA ligation which is a critical step in all repair pathways, appears to play a critical role in their resistance (***Nicolas et al., in press***). It is likely involved in BER but could potentially also be used for the ligation step of other repair pathways. Two other RNA ligases (FUN_027119/FUN_021735) predicted as HGT, were found OE only in the core response to radiation and with high TPMs (TPMs>100, 2.5h post-radiation, ***Supplementary Table S5***). These RNA ligases may play a role in nucleic acid repair as it was suggested for the RNA ligase DraRnl, upregulated in *Deinococcus radiodurans* after radiation [149]. HGT was suggested to have played an important role in the evolution of 2OG oxygenases in bacteria which were found to have a wide and uneven distribution within bacteria [97]. In our study, 20 amongst 49 genes characterized with a Pfam of this 2OG-Fe(II) oxygenase were characterized as HGTs, although some might have been missed as HGTs since their best blast hit were genes from fungi or bacteria. This family may play a role in the resistance to extreme conditions in bdelloid rotifers.

### 5. *A. vaga* exhibits similar molecular responses to both high- and low-LET radiation

This study allowed us to compare the biological and molecular responses to low- (X-rays) versus high-LET (Fe-ions) radiations, known to induce different type of DNA damages (base lesions, abasic sites, SSBs, DSBs; [57–60]). Moreover, damages induced by low LET are more homogeneously distributed in the cells and along the DNA than when exposed with Fe-ions which induce concentrated clusters of DNA damages [19, 64]. The harmful impact of high-LET radiation on the reproduction activity of *A. vaga* confirmed what was already observed by [19], as Fe-ions irradiation was associated with a severe sterilization of irradiated population (9±5% of fertile individuals post 500 Gy of Fe-ions vs 92±7% after 500 Gy X-ray radiation).

Our findings revealed that the molecular actors involved in the radiation response of *A. vaga* seem to be independent of radiation type. The overall transcriptomic response observed was strikingly similar between high- and low-LET-radiation, with log2foldchange and TPM values for specific genes (antioxidant, DNA repair-associated genes, HSPs) falling within the same range (***Figs 4B***, ***5B***) and 648 genes found OE in the core response to radiation (***Figs 3A***, ***3B***). We found some genes coding for antioxidants which seem specific to Fe-ions radiation, however these genes did not reach high TPMs or log2fc values.

We noticed one small difference: a time shift in the transcriptomic response between X-ray and Fe-exposed samples. While the major pattern of differentially expressed genes for *A. vaga* individuals exposed to 500 Gy occurred 2.5h post-rehydration, the response 8h post-Fe-ions irradiation was consistently larger than the response reported 8h post-X-ray radiation (e.g., ***Figs 1C***, ***2D***). This shift could be due to a delayed response in Fe-exposed animals, which are expected to deal with more complex damages, resulting in slower recovery and a postponed transcriptomic response compared to those irradiated with X-rays. The delayed reactivation of rehydrated *A. vaga* individuals might also be attributed to a longer desiccation period (5 days) during Fe-ion exposure (because of transport reasons, see Methods section) compared to X-ray irradiated samples (2 days desiccation). Extended desiccation periods were indeed correlated with a more prolonged recovery time in comparison to shorter periods (B. Hespeels, pers. comm.).

We hypothesize that the similarity in the transcriptomic responses following high- and low-LET radiation is because organisms are never exposed on earth to IR. Consequently, a specific response to the more complex damage caused by high-LET radiation has not been selected for. *A. vaga* can probably use the same actors to face damages due to desiccation or IR, even if high-LET radiation induces more clustered DNA damages than desiccation or low-LET radiation. This potentially less-adapted response to high-LET radiation might account for the reduced fertility observed following Fe-ion irradiation.

The difference in fertility rates observed with high-LET radiation triggers the question of how the germline cells deal with damages from radiation and if the mechanisms differ compared to somatic cells. We found in this study that most actors being OE post IR and desiccation belonged to NHEJ and BER pathways. NHEJ is known to be less faithful than HR when repairing DSBs and could cause some genetic modifications. Since bdelloid rotifers are known to be eutelic with a fixed number of somatic cells once in the adult stage, they might be able to survive even if a few genetic modifications remain in their somatic genomes. However, cellular division and DNA replication were shown to still occur in the germline of *A. vaga* [23]. Therefore, the production of viable offspring requires the complete and faithful repair of damaged germline nuclei. Since 72% of the released eggs were unable to successfully complete embryological development after 500 Gy of Fe-ions, we hypothesized that the DNA repair process acting in the germinal cells of bdelloids was overtaken by the complexity of damage and was associated with structural genomic change leading ultimately to the stop of embryological development. Transcriptomes used in this study represent mostly somatic cells (oocytes being a small proportion of the rotifer’s body), and we therefore cannot conclude on the DNA repair pathways acting in the germline of *A. vaga*. Future studies should establish alternative methods (e.g., single cell sequencing, in situ hybridization) allowing the investigation of the DNA repair mechanisms in developing oocytes after radiation.

## Conclusions

The common transcriptomic response found in the bdelloid rotifer *Adineta vaga* after desiccation and radiation (summarized on ***Fig 7*** in pink) provides evidence that *A. vaga* resistance to high doses of ionizing radiation (IR) is likely derived from an evolutionary adaptation to anhydrobiosis, a stress-resistant state they routinely encounter in their semi-terrestrial environments, consistent with the hypothesis of ***Mattimore and Battista*** [9]. Indeed, essential genes involved in IR resistance seem to have already evolved to withstand desiccation, and have been co-opted to counteract radiation stress. Among those genes is the arsenal of proteome protection genes in *A. vaga*, with antioxidant genes and chaperones (HSPs) being constitutively highly expressed (***Fig 7*** in green). These genes, and those involved in proteolysis via ubiquitination over-expressed after both stresses, are critical actors in their resistance to desiccation and radiation as they preserve protein integrity, including those involved in DNA repair. If many DNA repair genes (NHEJ, BER) were found over-expressed both post rehydration and radiation, IR induced a higher over-expression of DNA repair genes (***Fig 7*** in blue) because it causes more DNA damage. In addition to these genes, desiccation resistance requires additional actors post-rehydration (***Fig 7*** in yellow) to deal with the impact of water loss such as the upregulation of more chaperones, HSPs, LEA proteins, antioxidants, glucose repressive and cytoskeleton proteins), and with metabolic reduction as seen in anhydrobiosis.

While our research has identified a common response of desiccation and IR resistance, it is necessary to investigate these resistances across different species with different resistance to radiation and desiccation, including desiccation sensitive species [8, 150]. We could then have a broader picture of how the resistance to the two stresses are connected. Finally, this study paves the road for reverse genetics experiments (e.g., knockouts) targeting promising candidate genes involved in DNA repair and coding for antioxidants but especially orphan genes with unknown functions (e.g., 58 genes found in the common response, ***Supplementary Table S9***) which seem critical and unique of bdelloid rotifer radiation and desiccation resistance.

## Methods

### Bdelloid rotifer cultures

Experiments were performed using isogenic *Adineta vaga* clones originated from a single individual from the laboratory of Matthew Meselson at Harvard University. The cultures have been kept hydrated at 21°C in 150 × 20 mm Petri dishes supplemented with natural spring water (Spa®) and fed weekly with sterile lettuce juice. *A. vaga* individuals used for the experiment aiming to evaluate the impact of 14 days of desiccation and 1.5h post rehydration were cultivated with the same condition in the exception of food source which was *Escherichia coli* 1655MG.

### Desiccation protocol

*A. vaga* individuals were dried following the optimized protocol previously described in [12]. Briefly, cultures were washed with 15 mL Spa® water the day before their collection. Individuals were detached from the petri dish with a cell scraper followed by a short round of vortex. Animals were transferred to a 15 mL Falcon tube for centrifugation. Pellets, from each petri dish, were pooled in a final tube. The concentrated pellet containing rotifers was resuspended in Spa® water to a concentration of 80.000 individuals per mL. Then, for each desiccation sample, 0.5 mL of liquid was placed in the center of a Petri dish containing 30 mL of 3 % Low Melting Point agarose (LMP agarose, Invitrogen). LMP agarose plates with hydrated individuals (approx. 40.000 per plate) were placed in a climatic chamber (WEKK 0028) for dehydration with the following parameters: (1) linear decrease in relative humidity (RH) from 70 % to 55 % for 17 h (Temperature: 21-23°C), (2) linear decrease in relative humidity from 55 % to 41 % for 1 h (Temperature: 21-23°C), and (3) maintenance at 41 % RH and 21°C for the desiccated period. Based on this protocol, it took approximately 37 hours after the start of the dehydration process to dry all *A. vaga* individuals.

### X-rays radiation

40.000 hydrated or one day desiccated bdelloids were irradiated with 225 kVp X-rays at a dose rate of ∼7.8 Gy/min (using a X-ray irradiator PXi X-RAD 225 XL) up to a final dose of 500 Gy. During irradiation, samples were maintained on a refrigerated water bag to mitigate heating due to soft X-rays.

### Fe-ions irradiation

Irradiations with high-LET 56Fe-ions were performed at the GSI Helmholtz Center for Heavy Ion Research in Darmstadt, Germany. The iron ion beam was accelerated up to 1 GeV/n in the SIS18 synchrotron and extracted in air in Cave A. Dosimetry and beam monitoring is described in [151]). The beam has a LET of 147 keV/micron in water, to be compared to around 2 keV/micron for the 225 kVp X-rays. The preparation of desiccated samples took place in Namur University according to the protocol described above. Samples containing 40.000 individuals were transported under refrigerated environment to GSI the day before the exposure. Two sets of 3 days old desiccated bdelloid rotifers were assembled inside a sample holder allowing the exposition of multiple samples during a single beam time. After the exposure, samples were stored at 4°C and retrieved to Belgium for analysis. Samples were rehydrated 5 days after their entry in desiccation.

### Survival and fertility

After desiccation or radiation, bdelloid individuals were cultivated (and potentially rehydrated using 15 mL Spa^®^ water) at 21°C for 48 h. Bdelloids were considered alive when they had fully recovered motility or when the mastax moved in contracted individuals. Living bdelloids were separated from dead ones by transferring the supernatant, containing the latter, to new plates. Subsequently, the survival rate was manually calculated by tallying the number of living and dead specimens in each Petri dish under a binocular microscope.

In instances where individual count surpassed approximately 1,000 individuals, extrapolation was employed. This was achieved by counting sixfold the number of living or dead specimens observed in a 2 μL sample drawn from a homogeneously mixed culture, with a final volume of 2– 5 mL.

The reproductive capacity was defined as the ability for each isolated *A. vaga* individual to lay eggs and to develop clonal populations after being desiccated and potentially irradiated. One day post rehydration, we tested the fertility isolating a minimum of 60 successfully rehydrated individuals per condition; each isolated female was deposited in a well of a 12-well petri plate. Each well was filled with 2 mL of Spa water and 50 μL of sterilized lettuce juice. After 30 days, wells were observed under a binocular stereoscope checking for: (1) the presence of a population (minimum 2 adults and 1 egg per well), (2) the presence of only eggs that did not hatch (+ eventually the single adult from the start defined as a sterile individual), and (3) the presence of only dead individual(s).

### RNA extraction and RNA sequencing

Total RNAs were extracted using the RNAqueous-4PCR Kit (Ambion, Austin) following manufacturer instructions. Minimum 500 ng RNA of each sample was delivered to Genomics core (UZLeuven, Belgium) or Genoscope (Paris, France). X-ray and Fe-ions irradiated RNA samples were treated as follows: RNA libraries were prepared using the TruSeq Stranded mRNA protocol (Illumina, San Diego, USA). Libraries were purified and evaluated using an Agilent 2100 bioanalyzer. RNA libraries were sequenced with an Illumina NextSeq 500 (Illumina, San Diego, CA USA) as paired-end 2 × 75 base pair read using the NextSeq version 2.5 mid or high-output 150 cycle kit (Illumina).

Samples studying the expression following 14 days of desiccation were treated as follows: total RNA was enriched in mRNA based on polyA tails, chemically fragmented and converted into single-stranded cDNA using random hexamer priming to then generate double-stranded cDNA. Next, paired-end libraries were prepared following the Illumina 222s protocol (Illumina DNA sample kit): briefly, fragments were end-repaired, 3’-adenylated, and ligated with Illumina adapters. DNA fragments ranging in size from 300 to 600 bp (including the adapters) were PCR-amplified using adapter-specific primers. Libraries were purified, quantified (Qubit Fluorometer; Life technologies), and library profiles were evaluated using an Agilent 2100 bioanalyzer. A paired-end flow cell of 101-bp reads was sequenced for each library on an Illumina HiSeq2000 platform.

The fastq files of each transcriptome are accessible on NCBI under the Bioproject PRJNA962496 under the biosamples SAMN34396008-SAMN34396072.

### Differential gene expression analysis (DGE)

The whole bioinformatic workflow described below was performed in the Galaxy Europe platform (usegalaxy.eu). Sequencing adapters and the first ten nucleotides were removed with Trim Galore (version 0.4.3.0) (http://www.bioinformatics.babraham.ac.uk/projects/trim_galore/). The quality of the transcriptomes was checked by using FASTQC evaluation software FastQC (version 0.67) (http://www.bioinformatics.babraham.ac.uk/projects/fastqc/) before and after trimming the adapters. Trimmed-reads were mapped onto the genome assembly 2020 of *A. vaga* [65] using RNA-Star (version 2.5.2b0) [152]. We transformed the gff file into a gtf file using gffread (version 2.2.1.1) [153] to use the gtf for the read mapping. A matrix of normalized counts per gene was assembled by htseq-count (version 0.6.1) [154].

We conducted the differential gene expression analysis (DGE) with all the data together, combining 15 control samples together, and combining 11 desiccated samples, using DESeq2 (version 2.11.39, [155, 156]) and EdgeR (version 3.28.0, [157, 158]) in the R statistical software (version 3.4.1) (http://www.R-project.org) (not performed in galaxy platform).

For subsequent analyses, genes were considered significantly differentially expressed in the treated samples compared to the control samples if they have both adjusted p-value < 0.01 for Deseq2 analyses and an adjusted false discovery rate (FDR) < 0.01 for EdgeR results, and absolute values of their log2fold-change ≥ 0.5 with both tools. Those with a log2fold-change ≥ 0.5 were more expressed in the treated samples than in the controls (categorized as over-expressed OE genes), whereas those with a log2fold-change ≤ -0.5 were less expressed in the treated samples than in the controls (under-expressed UE genes). In this study we used the values obtained with DESeq2 for the graphs and tables.

### DGE comparisons

We analyzed the transcriptomic responses of *A. vaga* individuals exposed to desiccation and/or high-/low-LET radiation compared to hydrated controls (n=15) (***Fig 1A***). In order to capture the transcriptomic pattern resulting from the dehydration response, we focused our analysis on *A. vaga* individuals entering desiccation after 48 hours (n=11). We also analyzed the transcriptomic response in *A. vaga* individuals rehydrated for 1.5h after being desiccated during 14 days (n=3) to study the impact on gene expression after a longer period of dryness.

To characterize the radiation response of *A. vaga* individuals to X-rays, hydrated and desiccated animals were exposed to 500 Gy. The transcriptomic response was investigated post-radiation exposure, and optional rehydration, at different time points: 2.5h, 8h and 24h (this latter timepoint was only investigated in previously desiccated rotifers) (n=4 for each time point). Comparison of radiation response in hydrated and rehydrated animals ensured the discrimination of radiation specific response from genes impacted by the dehydration/rehydration process. A similar experiment was performed by exposing only desiccated *A. vaga* individuals to 500 Gy Fe-ions radiation (n=4 for time points 2.5h, 8h, and 24h). This second kinetic was performed to compare the biological and transcriptomic response of irradiated *A. vaga* against low- or high-LET radiation.

### Transcripts per millions

Transcripts per million (TPMs) for each gene was computed with a custom Python script (given in the supplements).

### Gene annotations

Gene Ontology (GO) ids, Pfam domains, and KEGG ids were annotated using InterProScan [159] in Galaxy with *A. vaga* gene set [65]. We searched for putative protein functions by similarity searches using BLASTP in the non-redundant database (NRdb) using a threshold expect value E < 10^−5^. Annotated genes were validated by blast and *A. vaga* genome was manually curated to complete DNA repair pathways and antioxidant gene families.

### Horizontal gene transfer

Horizontal gene transfer (HGT) acquisition was evaluated based on Alienomics as previously calculated for the *A. vaga* gene set [65]. This approach only identifies putative horizontal gene transfers (HGTs) from non-metazoans (e.g., bacteria, plants or fungi).

### Gene Ontology enrichment analysis

GO enrichment analysis was performed in R, using the package “topGo” from Bioconductor [160], (script available under request).

### Pathway and heatmap

KO (KEGG Ontology) annotations were performed using bi-directional best hit with the KAAS annotation server [161]. Eleven supplemental organisms enriched the KAAS default gene dataset to increase the annotation quality. Based on KO numbers, pathway maps were designed using the KEGG pathway reconstruction tool [162–164].

DNA repair pathways (NHEJ, BER, NER, HR, MR) were represented based on the KEGG pathway map, respectively, map03450, map03410, map03420, map03440, map03430. The log_2_foldchanges of genes significantly differentially expressed (confirmed with EdgeR) obtained with Deseq2 were represented for each gene of the pathways, using the ComplexHeatmap package from Bioconductor [165].

## Declarations

### Ethics approval and consent to participate

Not applicable

### Consent for publication

Not applicable

### Availability of data and materials

All analyses described in this study are published as supplementary tables, and transcriptomic reads can be found on NCBI under the Bioproject PRJNA962496 under the biosamples SAMN34396008-SAMN34396072.

### Competing interests

The authors declare that they have no competing interests

### Funding

Project was supported by the European Space Agency (ESA) and the Belgian Federal Science Policy Office (BELSPO) in the framework of the PRODEX Program. Fe-ions irradiations were supported by ESA and GSI through IBER2017-2021 campaigns (Investigations into Biological Effects of Radiation). KVD was supported by the European Research Council (ERC) with the grant agreement 725998 (RHEA). S. Penninckx is funded by the Walloon Region (PROTHERWAL, grant n°7289).

### Author contributions

VCM: Transcriptomic analyses, figures, tables, core manuscript writing

LB: Preparation of samples for X-ray and Fe-ions irradiation, laboratory cultures, RNA extraction and purification

JB: preparation of samples for Fe-ion irradiation, laboratory cultures, survival and fertility assays ACH: Radiation experiment support

SP: Radiation experiment support

SR: Fe-ion irradiation experiment

UW: Fe-ion irradiation experiment

MD: Fe-ion irradiation experiment

EGJD: Support with preliminary transcriptomic analyses

BH: lab cultures, preparation of samples for X-ray and Fe-ion irradiation, survival and fertility rate, RNA extraction and purification, preliminary transcriptomic analyses, experimental design conception, manuscript writing, fund acquisitions

KVD: experimental design conception, follow-up on the results, manuscript writing, fund acquisitions, project supervisor

## Acknowledgement

The authors thank A. Houtain for his help writing Python script for TPM calculations, M. Da Rocha, C. Rancurel for their collaboration during data analysis of first transcriptomic data. C. Da Silva and O. Jaillon for their support during RNA sequencing of hydrated and desiccated samples.

## Supplementary Figures

**Fig S1.**
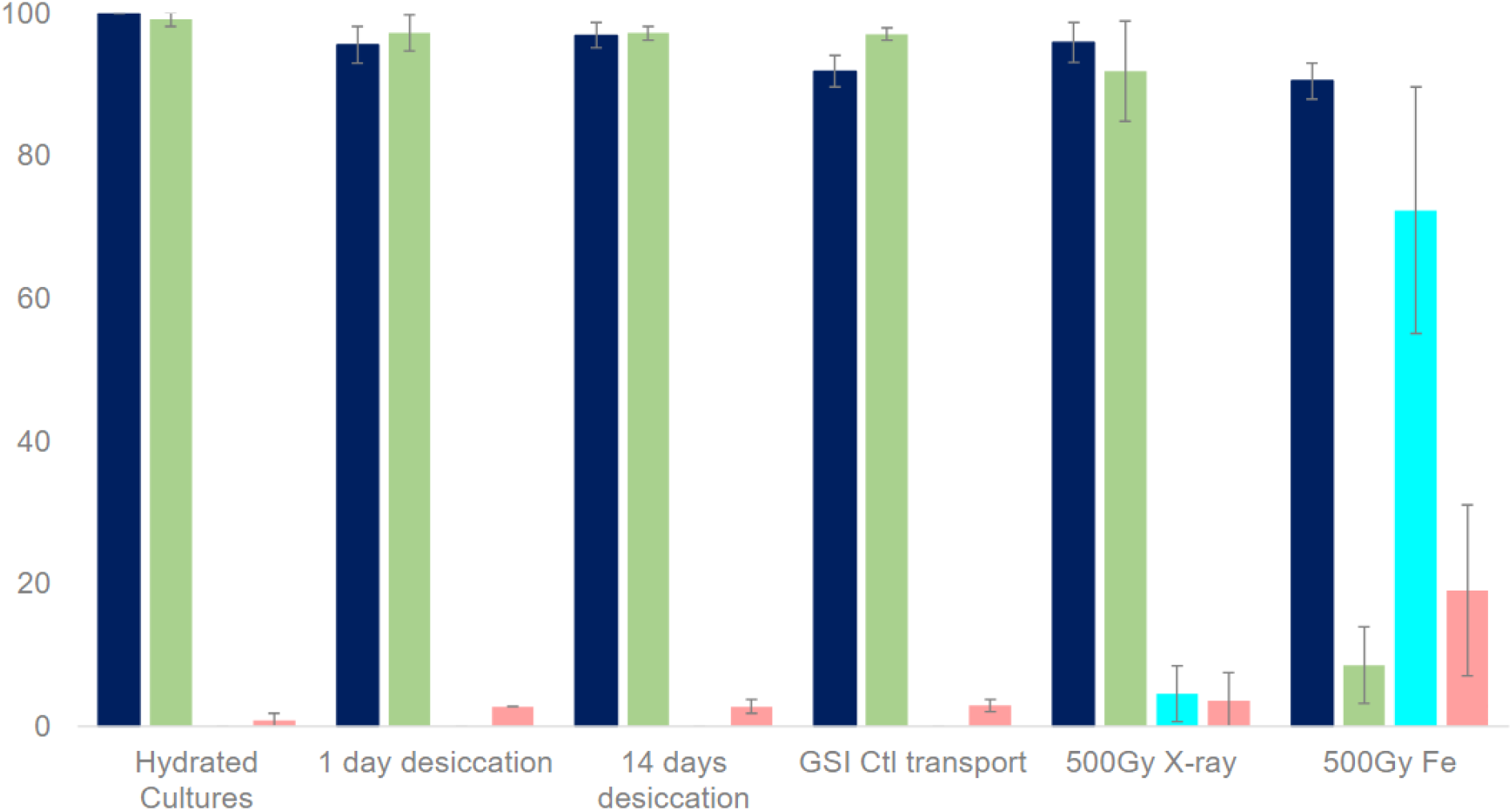
Survival and fertility rates of *A. vaga* individuals exposed to desiccation and radiation stress. Percentage of survival (dark blue) and fertility rate of *A. vaga* exposed to desiccation, rehydration post 14 days desiccation, and 500 Gy of X-rays (low-LET) or Fe-ions (high-LET). The fertility is represented by three different histograms: fraction of individuals able to produce viable offspring (green), fraction of individuals unable to produce viable offspring but only sterile egg(s) (blue), fraction of individuals unable to lay egg or with premature dead (red). Survival rate was evaluated 2 days post rehydration or post radiation. To ensure reliable results, survival data were obtained from at least three replicates. Samples exposed to 500 Gy of iron ions were transported from Belgium to GSI and then returned to the authors’ laboratory for analysis. To evaluate the impact of transportation on the samples, a control group labeled “Ctl transport” was also sent to GSI. This control group experienced similar conditions as the exposed samples but did not undergo radiation exposure.

**Fig S2.**
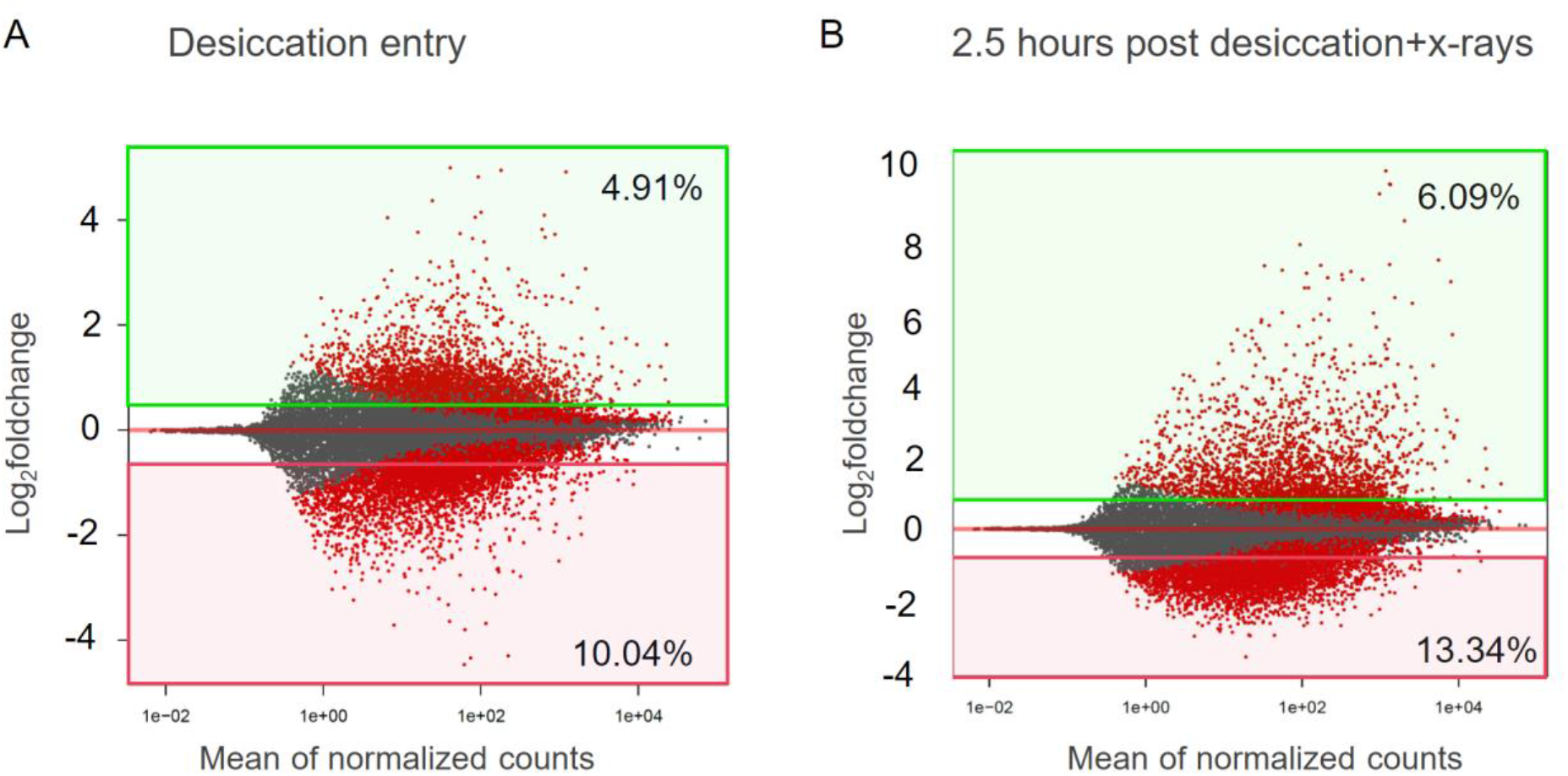
Volcano plots of differential genes over-expressed (OE) and under-expressed (UE) in A) *A. vaga* individuals entering desiccation, B) 2.5 hours post desiccation and x-rays radiation. The percentage of the genome over- and under-expressed are written. Although a higher percentage of genes are under-expressed in B) the log_2_foldchange values are bigger in OE than UE genes.

**Fig S3.**
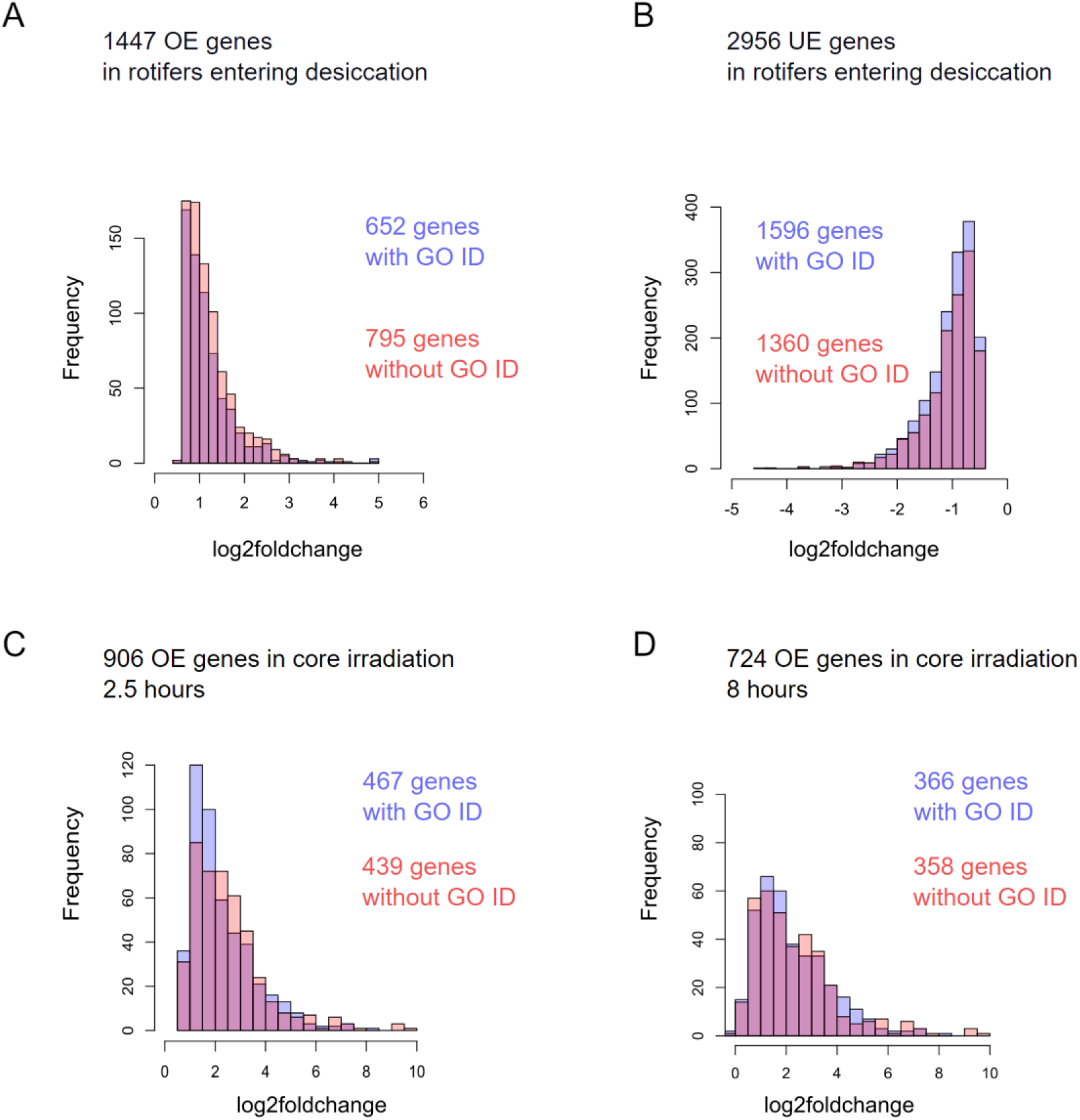
Number of genes with GO ids for a specific log_2_foldchange values identified as A) over-expressed (OE) genes in *A. vaga* rotifers entering desiccation, B) under-expressed (UE) genes in *A. vaga* rotifers entering desiccation, C) 906 OE genes in the core response to radiation 2.5 hours post irradiation, D) 724 OE genes in the core response to irradiation 8 hours post radiation.

**Fig S4.**
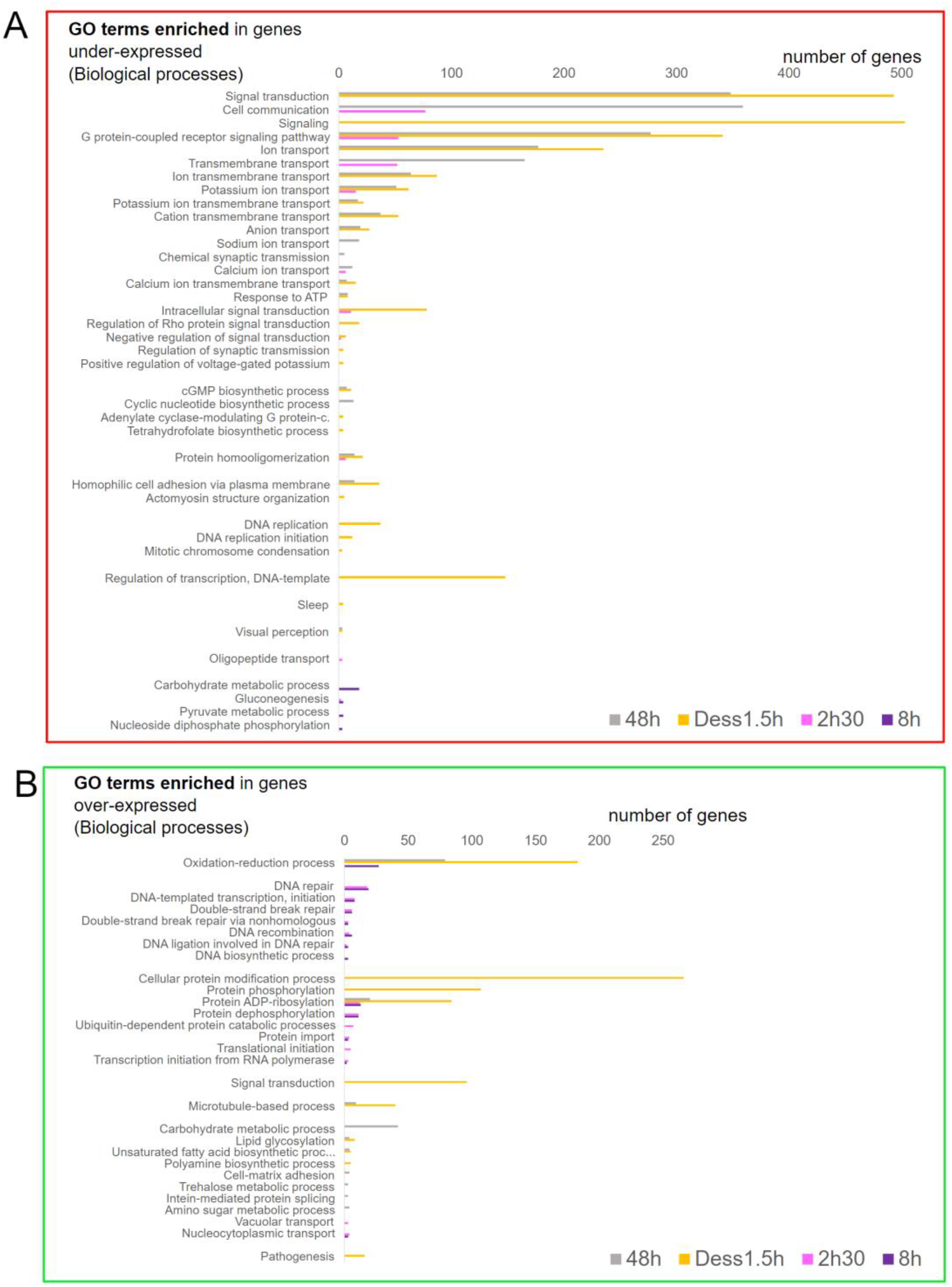
Gene Ontology enrichment analyses. GO biological processes significantly enriched (chi-square test p-value<0.05, min. 3 OE genes with the GO id) with genes being significantly A) under-expressed (framed in red, FDR<0.01 and log_2_foldchange < -0.5 in DESeq2 and EdgeR) and B) over-expressed (framed in green, FDR<0.01 and log_2_foldchange > 0.5 in DESeq2 and EdgeR) in *A. vaga* individuals entering desiccation (in gray) and 1.5 hours post-rehydration after 14 days of desiccation (in yellow), and in the core response to radiation 2.5h (pink) and 8h (purple) post radiation.

**Fig S5.**
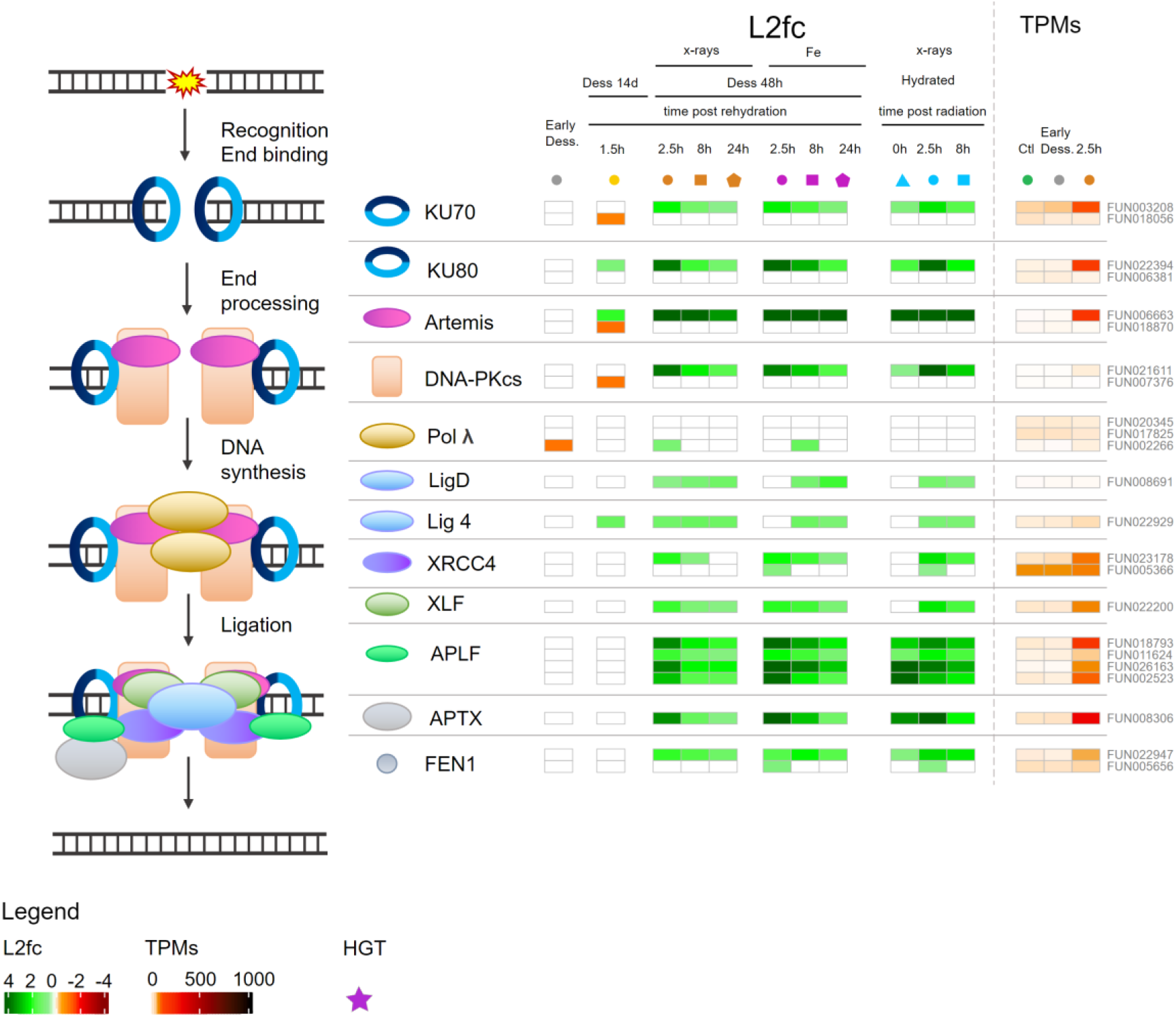
Expression of genes involved in Non-Homologous End-Joining (NHEJ) pathway differentially expressed post desiccation and/or radiation. The figure represents how the different actors act in this DNA repair pathway and in which order. They are represented by different symbols. How the level of expression of genes (gene ids written on the right side of the figure) coding for these actors change in the different conditions is represented by a heatmap of the log2foldchange. The color is not white when the gene is significantly (FDR < 0.01 with both DESeq2 and EdgeR) over-(log2foldchange>0.5 in green) or under-expressed (log2foldchange<- 0.5 in orange or red). The investigated conditions are: individuals entering desiccation (gray), after 14 days desiccation and 1.5 hours rehydration (yellow), after desiccation and x-rays radiation (orange), after desiccation and Fe radiation (purple), after x-ray radiation without desiccation (blue). The investigated time points are represented as follow: 0 hours (triangles), 1.5 or 2.5 hours (circles), 8 hours (squares), 24 hours (pentagons). Heatmap of transcripts per millions (TPMs) for each gene is represented on the right side and with a different code representing the level of expression. Horizontal transfer genes (HGT) are represented by the violet stars.

**Fig S6.**
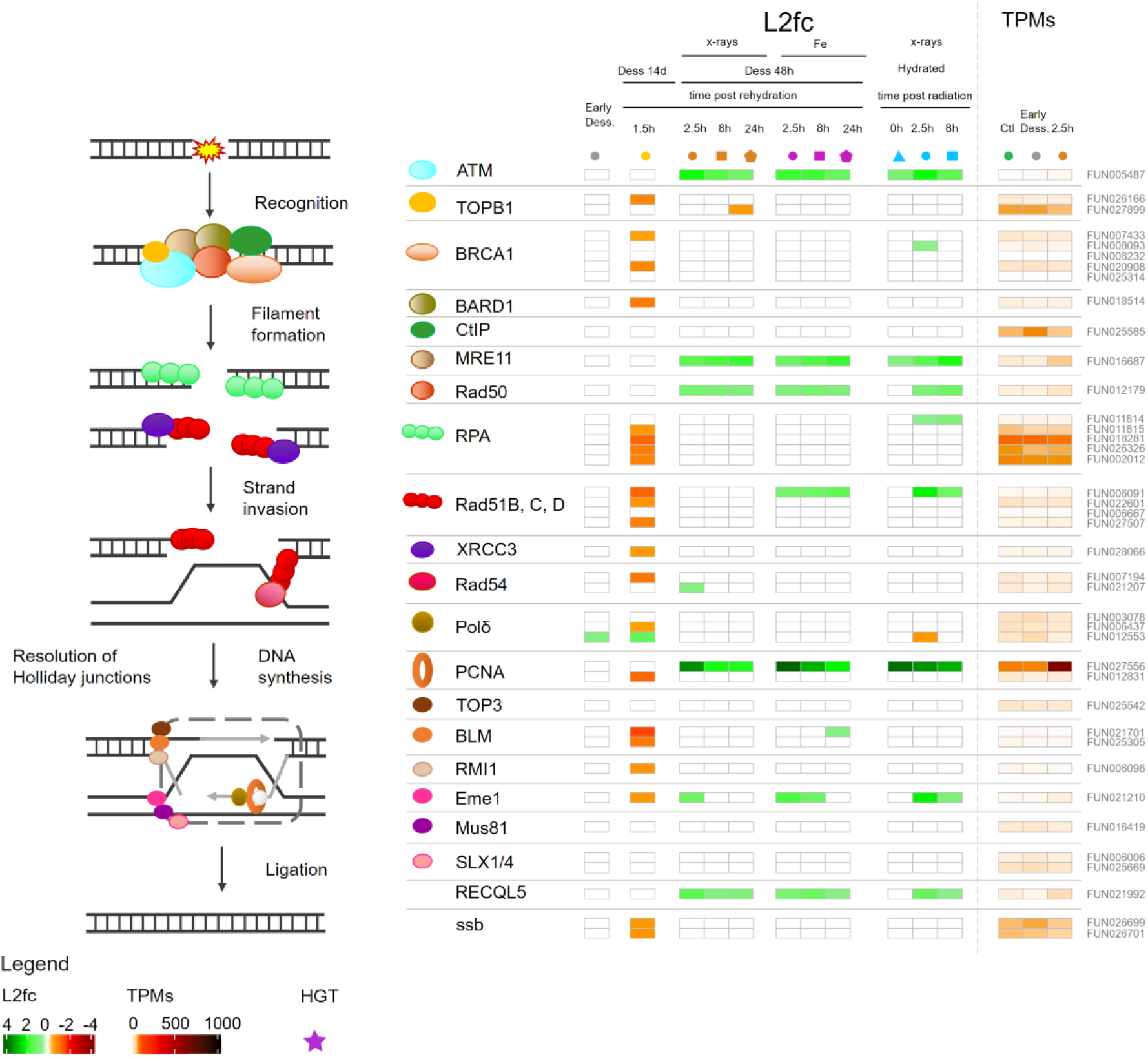
Expression of genes involved in Homologous Recombination (HR) pathway differentially expressed post desiccation and/or radiation. The figure represents how the different actors act in this DNA repair pathway and in which order. They are represented by different symbols. How the level of expression of genes (gene ids written on the right side of the figure) coding for these actors change in the different conditions is represented by a heatmap of the log2foldchange. The color is not white when the gene is significantly (FDR < 0.01 with both DESeq2 and EdgeR) over-(log2foldchange>0.5 in green) or under-expressed (log2foldchange<- 0.5 in orange or red). The investigated conditions are: individuals entering desiccation (gray), after 14 days desiccation and 1.5 hours rehydration (yellow), after desiccation and x-rays radiation (orange), after desiccation and Fe radiation (purple), after x-ray radiation without desiccation (blue). The investigated time points are represented as follow: 0 hours (triangles), 1.5 or 2.5 hours (circles), 8 hours (squares), 24 hours (pentagons). Heatmap of transcripts per millions (TPMs) for each gene is represented on the right side and with a different code representing the level of expression. Horizontal transfer genes (HGT) are represented by the violet stars.

**Fig S7.**
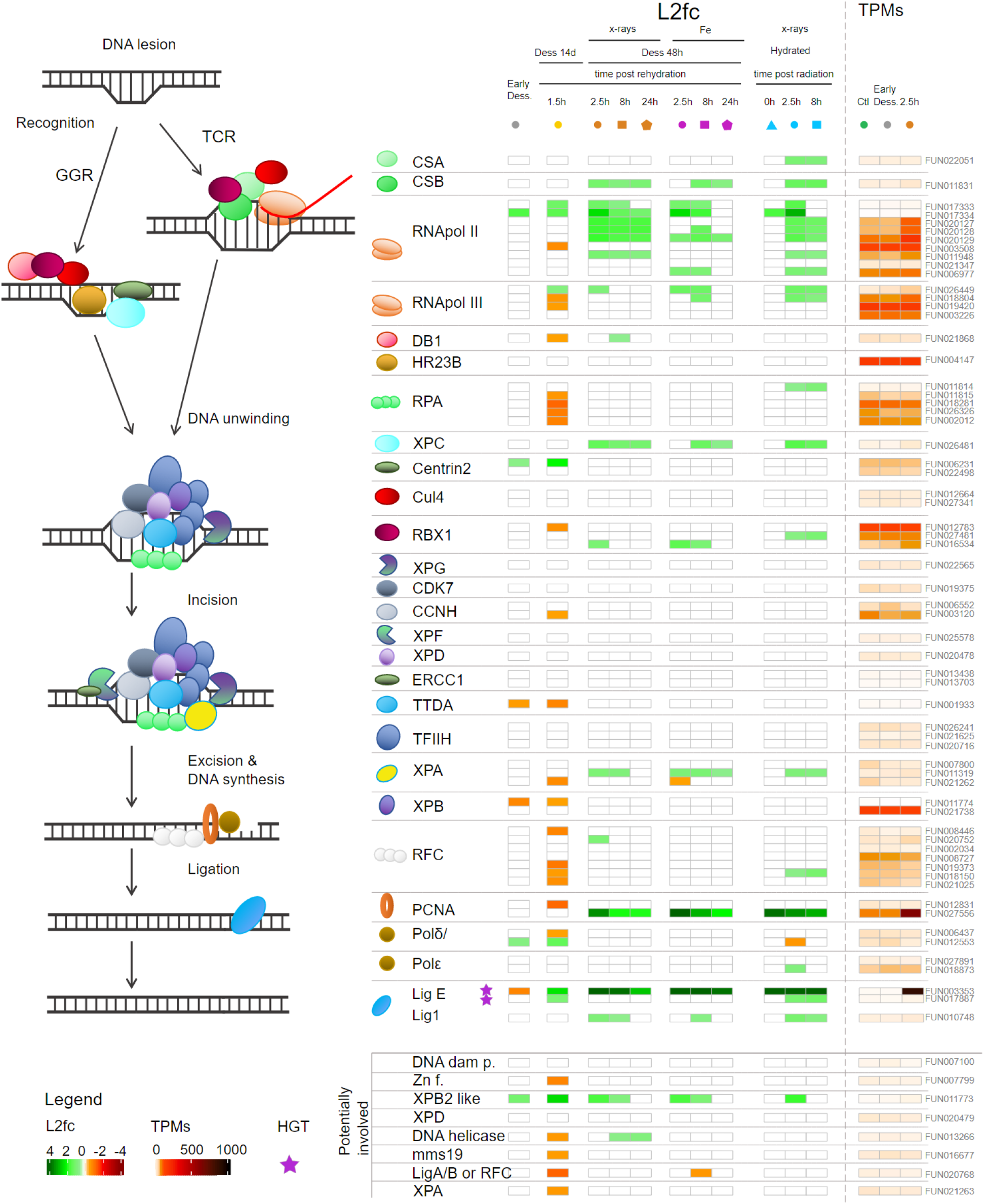
Expression of genes involved in Nucleotide excision repair (NER) pathway differentially expressed post desiccation and/or radiation. The figure represents how the different actors act in this DNA repair pathway and in which order. They are represented by different symbols. How the level of expression of genes (gene ids written on the right side of the figure) coding for these actors change in the different conditions is represented by a heatmap of the log2foldchange. The color is not white when the gene is significantly (FDR < 0.01 with both DESeq2 and EdgeR) over-(log2foldchange>0.5 in green) or under-expressed (log2foldchange<- 0.5 in orange or red). Genes potentially involved in this pathway are indicated in the bottom of the figure. The investigated conditions are: individuals entering desiccation (gray), after 14 days desiccation and 1.5 hours rehydration (yellow), after desiccation and x-rays radiation (orange), after desiccation and Fe radiation (purple), after x-ray radiation without desiccation (blue). The investigated time points are represented as follow: 0 hours (triangles), 1.5 or 2.5 hours (circles), 8 hours (squares), 24 hours (pentagons). Heatmap of transcripts per millions (TPMs) for each gene is represented on the right side and with a different code representing the level of expression. Horizontal transfer genes (HGT) are represented by the violet stars.

**Fig S8.**
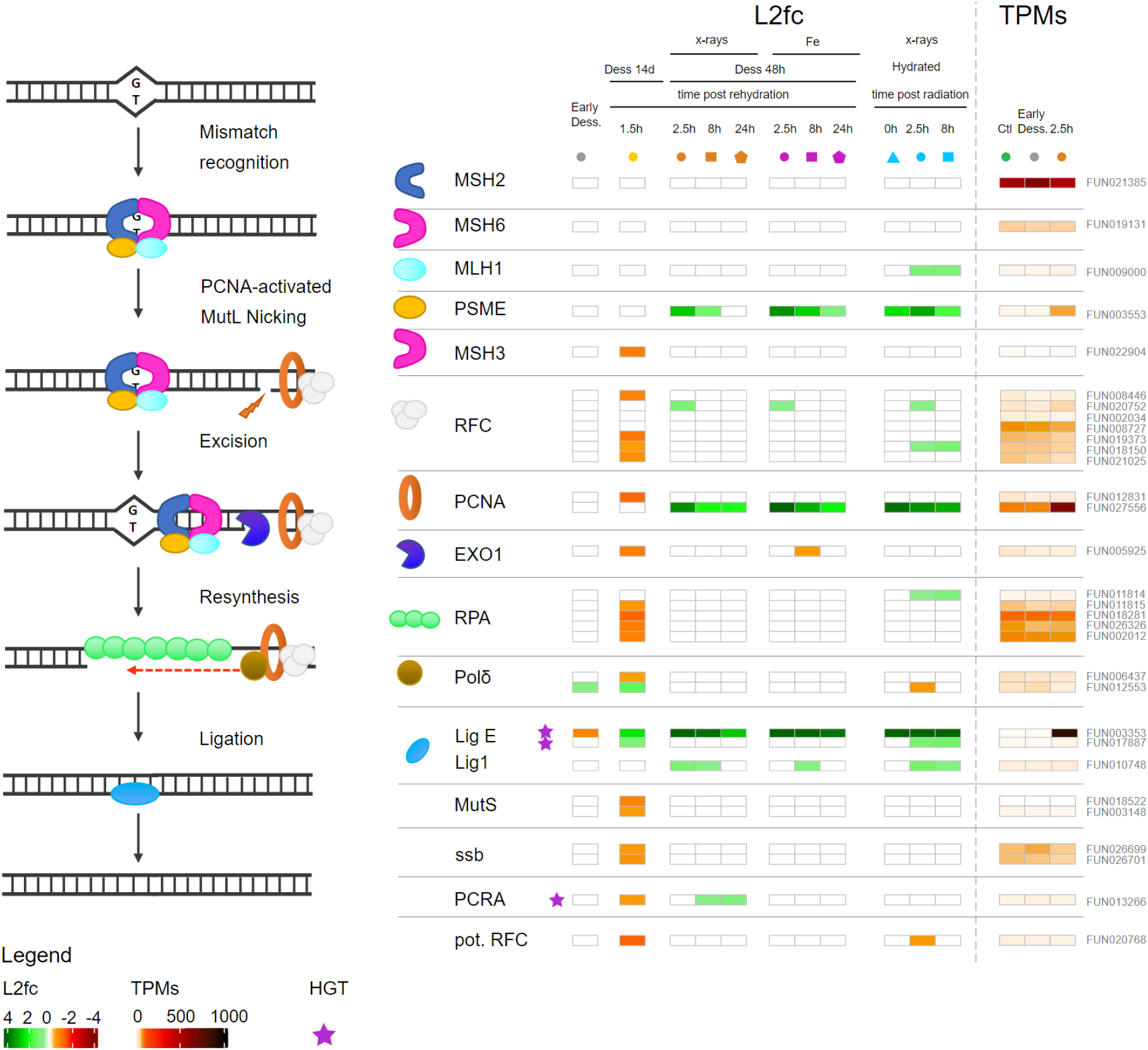
Expression of genes involved in Mismatch repair pathway (MR) differentially expressed post desiccation and/or radiation. The figure represents how the different actors act in this DNA repair pathway and in which order. They are represented by different symbols. How the level of expression of genes (gene ids written on the right side of the figure) coding for these actors change in the different conditions is represented by a heatmap of the log2foldchange. The color is not white when the gene is significantly (FDR < 0.01 with both DESeq2 and EdgeR) over-(log2foldchange>0.5 in green) or under-expressed (log2foldchange<-0.5 in orange or red). Genes potentially involved in this pathway are indicated in the bottom of the figure. The investigated conditions are: individuals entering desiccation (gray), after 14 days desiccation and 1.5 hours rehydration (yellow), after desiccation and x-rays radiation (orange), after desiccation and Fe radiation (purple), after x-ray radiation without desiccation (blue). The investigated time points are represented as follow: 0 hours (triangles), 1.5 or 2.5 hours (circles), 8 hours (squares), 24 hours (pentagons). Heatmap of transcripts per millions (TPMs) for each gene is represented on the right side and with a different code representing the level of expression. Horizontal transfer genes (HGT) are represented by the violet stars.

## Supplementary Tables

**Supplementary Table S1.** Genes within Adineta vaga genome. The columns give the gene ids from the gene assembly in Simion et al. (2020), the correspondence gene ids in the genome published by Simion et al. (2021), if the gene is annotated as HGT, the blast hits, the KEGG ids, GO ids, pfam domains, the results for all analyses with DESeq2 and EdgeR, and the TPM values for all the different analyzed conditions. The columns with the results from DESeq2 and EdgeR are explained in the second table sheet.

**Supplementary Table S2.** GO analysis of Under-Expressed genes in the core radiation response.

**Supplementary Table S3.** Genes found over-expressed in the core common response to radiation and desiccation. The columns give the gene ids, if the gene is annotated as HGT, its hypothesized function, its function acronym, pfam domains, best blast hits, the results for all analyses with DESeq2 and EdgeR, and the TPM values for all the different analyzed conditions. The columns with the results from DESeq2 and EdgeR are explained in the second table sheet of Supplementary Table S1.

**Supplementary Table S4.** Genes found over-expressed in the core rehydration response. The columns give the gene ids, if the gene is annotated as HGT, its hypothesized function, its function acronym, pfam domains, best blast hits, the results for all analyses with DESeq2 and EdgeR, and the TPM values for all the different analyzed conditions. The columns with the results from DESeq2 and EdgeR are explained in the second table sheet of Supplementary Table S1.

**Supplementary Table S5.** Genes found over-expressed in the core irradiation response. The columns give the gene ids, if the gene is annotated as HGT, its hypothesized function, its function acronym, pfam domains, best blast hits, the results for all analyses with DESeq2 and EdgeR, and the TPM values for all the different analyzed conditions. The columns with the results from DESeq2 and EdgeR are explained in the second table sheet of Supplementary Table S1.

**Supplementary Table S6.** List of genes coding for the defined antioxidant gene families. The columns give the gene ids, the antioxidant family, if the gene is annotated as HGT, the blast hits, the KEGG ids, GO ids, pfam domains, the results for all analyses with DESeq2 and EdgeR, and the TPM values for all the different analyzed conditions. The columns with the results from DESeq2 and EdgeR are explained in the second table sheet of Supplementary Table S1.

**Supplementary Table S7.** 34 Genes coding for antioxidants exclusively significantly over-expressed in *A. vaga* individuals 1.5 hours post rehydration after 14 days of desiccation. The columns give the gene ids, if the gene is annotated as HGT, the antioxidant family, the pfam domains, the results for all analyses with DESeq2 and EdgeR for all the different analyzed conditions. The columns with the results from DESeq2 and EdgeR are explained in the second table sheet of Supplementary Table S1.

**Supplementary Table S8.** List of genes coding for proteins involved in DNA repair. The columns give the gene ids, if the gene is annotated as HGT, the DNA repair pathway in which the gene is involved, its hypothesized functions (acronym and description) based on blast hits, on the KEGG ids, GO ids, and on pfam domains, the results for all analyses with DESeq2 and EdgeR, and the TPM values for all the different analyzed conditions. The columns with the results from DESeq2 and EdgeR are explained in the second table sheet of Supplementary Table S1.

**Supplementary Table S9.** List of genes with unknown functions but high log_2_foldchange and TPMs post radiation and/or desiccation. The columns give the gene ids, if the gene is annotated as HGT, its hypothesized function, its function acronym, pfam domains, best blast hits, the results for all analyses with DESeq2 and EdgeR, and the TPM values for all the different analyzed conditions. The columns with the results from DESeq2 and EdgeR are explained in the second table sheet of Supplementary Table S1.

## Notes

### Competing Interest Statement

The authors have declared no competing interest.

### Summary of Updates

Discussion section updated to clarify Figures revised

